# Polyethylene valorization by combined chemical catalysis with bioconversion by plastic-enriched microbial consortia

**DOI:** 10.1101/2022.12.20.521199

**Authors:** Gwendolyn J. Gregory, Cong Wang, Sunitha Sadula, Sam Koval, Raul Lobo, Dionisios G. Vlachos, E. Terry Papoutsakis

**Affiliations:** Department of Chemical and Biomolecular Engineering, University of Delaware, Newark, DE; Delaware Biotechnology Institute, University of Delaware, Newark, DE; Center for Plastics Innovation, University of Delaware, Newark, DE; Catalysis Center for Energy Innovation, 221 Academy St., Newark, DE; Department of Biological Sciences, University of Delaware, Newark, DE

**Keywords:** plastics, biodegradation, polyolefins, upcycling, circular economy, biocatalysis, environmental consortia

## Abstract

There are few reports of microbial deconstruction or functionalization of the recalcitrant backbone of polyolefins. However, microbes can utilize polyolefin deconstruction products, including n-alkanes. Here, we combined chemical catalysis with bioconversion to valorize polyethylene (PE) deconstruction products. High-density PE (HDPE) was deconstructed via hydrogenolysis over a ruthenium on carbon catalyst. The resulting *n*-alkane mixture (C_4_-C_35_) was utilized as a feedstock for microbial consortia derived from soil from local recycling plants. We found two consortia that utilized the PE-deconstruction product mix as a sole carbon source. We adapted the consortia on a commercially-available *n*-alkane mix to reduce the number of species present and enrich for enhanced alkane utilization. Both resulting enriched consortia utilized the PE-deconstruction product mix more effectively than the original (parent) consortia. The predominant metabolite produced by both enriched consortia was a C_16_-C_16_ wax ester. Wax esters have considerable industrial value, with the longer chain lengths (C_32_-C_36_) having the highest value. We identified two *Rhodococcus aetherivorans* strains that grow well on C_24_, indicating this species is important for the functionalization of long-chain alkanes. This work demonstrates that enriched consortia from plastic-enriched environments can be combined with chemical catalysis to valorize polyethylene.

**Synopsis:** Chemical catalysis can be used to deconstruct polyethylene waste material to produce a mixture of alkanes. Enriched environmental microbial consortia can valorize these polyethylene deconstruction products via functionalization that preserves the alkane chain length thus minimizing CO_2_ production.

## Introduction

The accumulation of plastic waste in the environment is of urgent concern. An estimated 400 million metric tons of plastic are produced annually, approximately 60% of which are polyolefins, such as polyethylene (PE) and polypropylene^1, 2^. Given the low fraction of plastics recycling^2^ and the limited utility of polyolefin mechanical recycling and incineration, there is a pressing need to develop innovative polyolefin recycling via biological or chemical processes. In fact, the degradation of commercial polyethylene by a wide variety of bacterial genera was recently reported (see reviews^1, 3^). Very few enzymes, however, have been identified that can functionalize PE^4–6^, with no reported enzymes that can cleave the recalcitrant C-C backbone of PE.

Progress has been made in recycling via PE catalytic deconstruction, which produces linear or branched alkanes with size distribution in the diesel and wax/lubricant base-oil range^7–12^. Metal-catalyzed hydrogenolysis catalysts have good reactivity and selectivity at low temperatures. Ruthenium on tungstated zirconia (Ru-WZr) catalyst, for example, is highly active on low-density polyethylene (LDPE) at mild conditions and significantly repressed the generation of methane, a low-value byproduct typically produced by Ru-supported catalysts. The product is a mixture of liquid linear alkanes with a chain-length distribution of C_4_ to C_35_^13^.

While the identification of enzymes that can deconstruct or functionalize the recalcitrant backbone of polyolefins has been challenging, it is well known that microbes can utilize many of the deconstruction products from catalytic polyolefin breakdown, including linear alkanes^14–16^. Microbial conversion of PE deconstruction products with bacterial isolates has been reported previously^17–19^ and microbial degradation of alkanes has been investigated for years in the context of bioremediation of oil-contaminated sites^20–23^. Several species of bacteria can metabolize a broad range of alkanes. *Acinetobacter sp.* DSM17874, for example, has an *n*-alkane substrate range of C_10_-C_40^24^_. However, microbial consortia utilize a mixture of hydrocarbons better than a single species due to a wider range of metabolic pathways and enzymes present in the consortia^25,26^. Studies of alkane degradation using microbial consortia have either developed artificial consortia for hydrocarbon degradation or used reconstructed environmental consortia^25, 27–29^. One such study utilized an enrichment procedure to reduce the complexity of the consortium and select for highly efficient hydrocarbon degraders^25^.

Here, we integrate chemical catalysis with microbial conversion to functionalize and valorize PE-deconstruction products (Figure 1). We screened environmental soil-derived microbial consortia collected from biofilms formed on PE sheets for utilization of alkane mixes (C_4_-C_35_) produced via PE hydrogenolysis. As linear alkanes are found in PE sheets^30^, these consortia have been enriched for microbes utilizing linear alkane substrates. We quantified the degradation and conversion of the PE-deconstruction products mix and identified metabolites produced by the consortia for polymer plastics valorization. The predominant metabolite produced by enriched consortia was a C_16_-C_16_ wax ester. We demonstrate the tunability of microbial consortia to PE deconstruction products and move toward utilizing mixed plastics waste as feedstock to produce valuable compounds.

**Figure 1.**
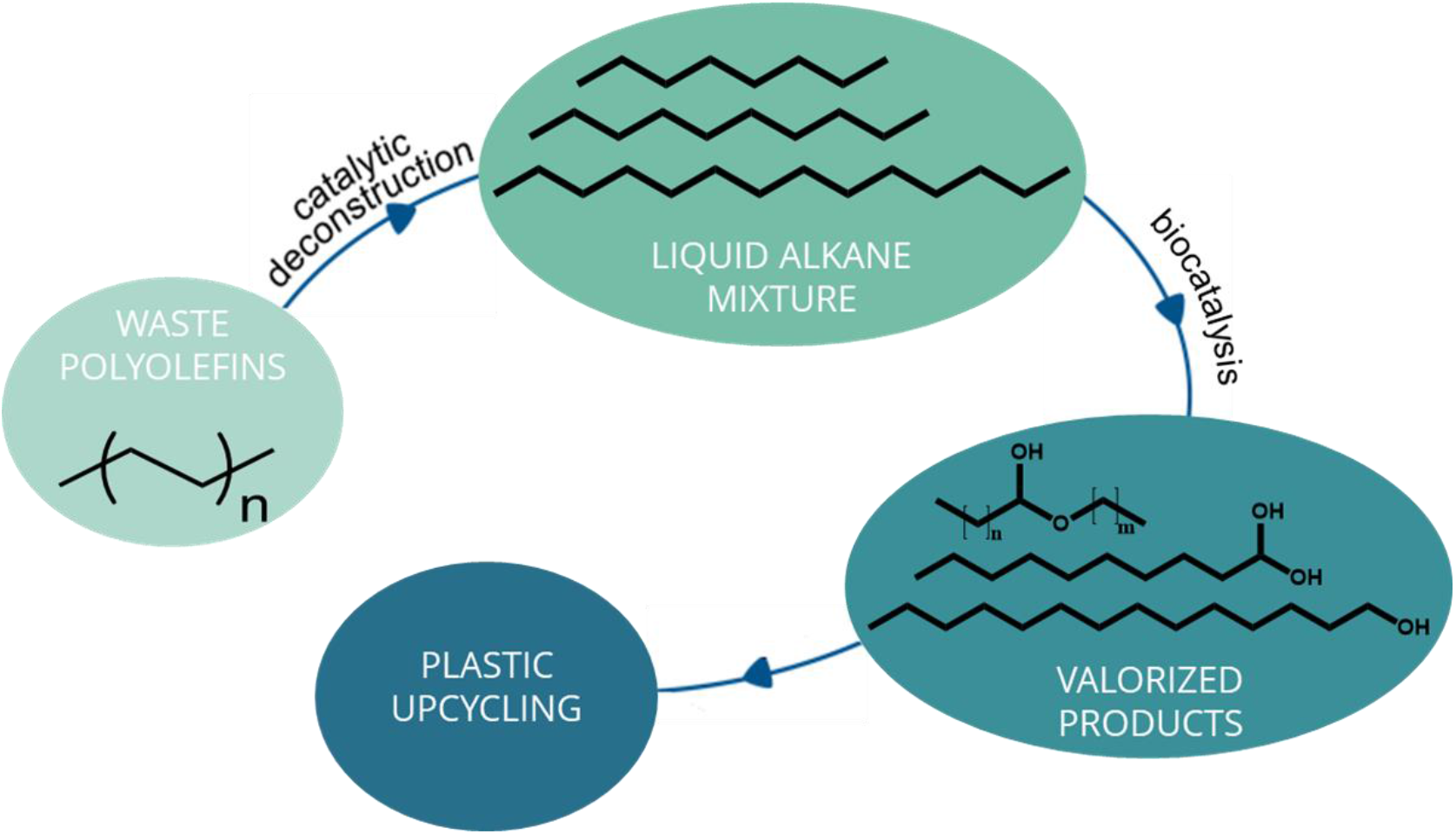
Upcycyling of waste polyolefins using chemical catalysis combined with bioconversion. PE is deconstructed over a Ru/C catalyst to produce a C_4_-C_35_ liquid alkane mixture. This mixture is used as a feedstock for microbial consortia that have been enriched for alkane utilization. The microbial consortia functionalize the linear alkane mixture, resulting in valorized products.

## Methods

### Environmental sampling and plastic enrichment

Soil samples were taken from the yard of a local recycling plant (Recycling Management Resources, Wilmington, DE) where water used to wash plastic waste was discharged and/or where plastic debris had accumulated. Samples were also collected from the sludge at the bottom of a cooling tank used to cool extruded recycled plastics (Eco Plastic Products, Wilmington, DE). PE sheets were sterilized with 70% ethanol and dried in sterile air. Approximately 1 g of soil/sludge from each sample site was incubated in duplicate in glass test tubes with 1 cm x 2 cm x 0.25 mm polyethylene sheets (Goodfellow, Coraopolis, PA) at 30°C with shaking (200 RPM) for 7 months in minimal medium (MM) (0.05% yeast extract, 0.2% ammonium sulfate, and 1% trace elements (0.1% FeSO_4_·7H_2_O, 0.1% MgSO_4_·7H_2_O, 0.01% CuSO_4_·5H_2_O, 0.01% MnSO_4_·5H_2_O, and 0.01% ZnSO_4_·7H_2_O) in 10 mM phosphate buffer (pH 7.0))^31^. The microbial biofilm that formed on each PE sheet was collected by incubating with 0.9% NaCl solution overnight while shaking at 30°C. The sheets were vortexed in the solution to create a suspension, from which glycerol (20%) or DMSO (7%) stocks were made and stored at −80° C for later use. Initial growth tests were performed with selected biofilm communities, or consortia, to determine alkane utilization. Growth tests were performed in glass test tubes with 10 mL of MM with no yeast extract, supplemented with 35 μL of hexadecane as the sole carbon source. The cultures were inoculated with 100 μL of a frozen DMSO stock of each of seven consortia (1-7) and grown at 37°C for six days. Two consortia (1 and 2) that grew well on a model alkane hexadecane were chosen for further analysis.

### Catalyst preparation, polyethylene hydrogenolysis and liquid product collection and analysis

The 5-wt% Ru on carbon catalyst (Ru/C) was prepared using incipient wetness impregnation^13, 32, 33^. Ruthenium (III) nitrosyl nitrate solution (Ru(NO)(NO_3_)_x_(OH)_3_, Aldrich, 1.5 wt% Ru) was impregnated on the carbon support (Vulcan), dried at 110°C for 12 hr and reduced in a 100-sccm, 10% H_2_/He flow at 250°C for 2 hr. Polyethylene hydrogenolysis was performed as described elsewhere^13, 32–34^. Briefly, 2 g of low-density polyethylene (HDPE, Mw ~76 kDa) was mixed with 50 mg catalyst and transferred into a 50 mL stainless-steel Parr reactor with a 700 μL stir bar. The sealed reactor was purged with H_2_ completely and was charged to 30 bar H_2_ at ambient temperature. The reactor was heated up to 250°C in 20 min, maintained at 250°C for 3 hr under stirring, and quenched in a water/ice bath to below 10°C before collecting liquid products for microbial experimentation. The liquid products from the HDPE hydrogenolysis, typically consisting of C_4_ – C_35_ alkanes, hereafter referred to as PE-deconstruction product mix, were collected by syringe filtration to remove the catalyst. Quantification of the product distribution was performed using a gas chromatograph (GC, Agilent 7890A) equipped with an HP-1 column and a flame ionization detector (FID) using dilute solutions^13, 33^. The liquid product was dissolved into a calculated volume of a n-octacosane in dichloromethane solution (n-C28/CH_2_Cl_2_, 20mg C28 external standard in 20mL CH_2_Cl_2_). GC responses were calibrated using standard calibration mixtures.

### Alkane utilization analyses

We tested two consortia (1 and 2) that were able to utilize linear alkanes as a sole carbon source. Two precultures were inoculated with 350 μL biofilm glycerol stock in 10 mL of Bushnell-Haas medium (HiMedia Labs, Lincoln University, PA) (MgSO_4_ 2 g/L, CaCl_2_ 0.02 g/L, KH_2_PO_4_ 1 g/L, K_2_HPO_4_ 1 g/L, NH_4_NO_3_ 1 g/L, FeCl_3_ 0.05 g/L) in glass test tubes with 30 μL of the PE-deconstruction product mix. Precultures were grown for 3 days at 30°C with aeration (200 RPM) and then stored at −80°C in 7% DMSO for use as inocula for two alkane degradation experiments, 1 and 2 (Exp1 and Exp2). To assess variability after our preculture procedure, we subcultured one of the original preculture DMSO stocks for 3 days and used this as the inoculum for experiment 3 (Exp3). For the alkane degradation assay, 350 μL of each preculture DMSO stock was inoculated into 10 mL of Bushnell-Haas medium with 30 μL of the PE-deconstruction product mix as the sole carbon source. Two independent cultures were tested in each experiment along with two negative controls containing just media and the PE-deconstruction product mix. After 14 days of growth, the remaining alkanes were extracted from the entire culture with 3 mL of dichloromethane (DCM) with an octacosane internal standard. The organic layer was collected and the extraction as repeated on the aqueous layer with 2 mL of DCM with an octocosane internal standard. The organic layer was collected and pooled. The residual concentration (mol %) was determined by GC-FID as described above. Percent conversion was calculated by dividing the percentage of recovered alkanes in the sample by the percentage remaining in the negative control and subtracting this from 100. The results of four independent cultures (two each from Exp1 and Exp2) were used to calculate average percent conversion and standard error of the mean (SEM). Individual experiments are the average of two independent cultures.

### Enrichment of consortia on liquid alkane mixture

DMSO stocks of consortia 1 and 2 (330 μL) were inoculated into 30 mL of Bushnell-Haas medium with 105 μL of a liquid alkane standards mixture (50 μg/mL of each even alkane from C_10_-C_40_ in hexane) (Restek, Bellefonte, PA) in 125 mL baffled Erlenmeyer flasks. The cultures were incubated at 37°C with shaking at 220 RPM for one week. Two mL of each culture were then passaged into fresh medium with 105 μL of the liquid alkane mixture and grown for two weeks with the same growth conditions. The cultures were passaged as before and grown for an additional four weeks. DMSO stocks were made of the entire culture and stored at −80°C for use as inocula for two alkane degradation experiments (Exp1 and Exp2). Alkane degradation assays were set up as described above. 10 mL of Bushnell-Haas medium with 30 μL of the PE-deconstruction product mix as the sole carbon source were inoculated with 350 μL of DMSO stocks of the enriched consortia 1 or 2. Cultures were grown for 14 days at 30°C with aeration. The remaining alkanes were extracted from the entire culture using 5 mL of DCM with an octacosane internal standard, as described above. Percent conversion was calculated as described above. The results of four independent cultures (two each from Exp1 and Exp2) were used to calculate average percent conversion and SEM. Statistics were calculated using an unpaired Student’s T-test with two tails.

### Metabolite identification and quantification

DCM extractions used to quantify alkane degradation were also used for the initial identification of intermediate metabolites produced from hexadecane. Using GC-MS, the major metabolites after growth on Bushnell-Haas medium were identified as hexadecanol, hexadecanoic acid, hexadecyl hexadecanoate and tetradecyl hexadecanoate. Since wax esters are accumulated under nitrogen-limiting conditions, assays to quantify the production of wax esters were performed using Nitrogen Limited Minimal Medium (NLMM) (0.01% ammonium sulfate, and 1% trace elements (0.1% FeSO_4_·7H_2_O, 0.1% MgSO_4_·7H_2_O, 0.01% CuSO_4_·5H_2_O, 0.01% MnSO_4_·5H_2_O, and 0.01% ZnSO_4_·7H_2_O) in 10 mM phosphate buffer (pH 7.0)). One mL of DMSO stock of E1 or E2 was inoculated into 30 mL NLMM in a 125 mL Erlenmeyer flask with 150 μL of hexadecane as the sole carbon source and incubated for one week at 30°C. Ten mL of the preculture was used to inoculate 10 mL of fresh NLMM in 125 mL flasks with 100 μL of hexadecane as the sole carbon source. The cultures were grown at 30°C with aeration for one week and then the metabolites were extracted from the 20 mL culture using the Bligh & Dyer chloroform:methanol:water extraction method (1:2:0.8 v/v/v)^35^, incubated with shaking for 2 hours, followed by the addition of chloroform:water (1:1 v/v). The extractions were stored at 4°C overnight and then centrifuged at 7000 g at 4°C for 10 minutes. The organic phase was collected in glass tubes then dried completely under nitrogen stream at 65°C. The solids were resuspended in 2 mL of chloroform with an octacosane internal standard. The metabolites were analyzed using a GC-FID (Agilent 7890A, HP-1 column) using octacosane as an internal standard. The injection temperature was 300°C. The column temperature program was: 40 °C (5 min), ramp at the rate of 20 °C/min to 300 °C and hold at 300 °C (5 min). The detection temperature was 300 °C. The products were identified by a GC (Agilent 7890B) mass spectrometer (MS) (Agilent 5977A with a triple-axis detector) equipped with a DB-5 column, high-resolution MS with liquid injection field desorption ionization. Three independent experiments were performed for each consortium with two replicate cultures each. Cell dry weight (CDW) was determined for each enriched consortium by filtering 20 mL of culture through a 0.22 μM polyethersulfone filter, washing gently with water and drying to constant weight.

### 16S sequencing of consortia and bioinformatics

To compare microbial community compositions, metagenomic DNA was extracted using Qiagen DNeasy Blood and Tissue Kit (Cat no. 69504) (Qiagen, Germany), according to the manufacturer’s protocol, from 350 μL of DMSO stock of each consortium preculture (Day 0) and samples collected after growth on the alkane mix for 14 days (Day 14). Partial 16S rRNA (V4, fourth hypervariable region of 16S gene) was amplified using 30 ng template DNA. Primers 515F and 806R with Illumina adapters and dual indices (8 bp)^36^ were used to amplify the V4 region. Pooled PCR products were sequenced on the MiSeq using v2 2×250 base pair kit (Illumina, San Diego, CA). Sequences were processed in Mothur v. 1.36.1, as previously described^36^. Following demultiplexing, sequences were clustered at 97% similarity. Relative abundances were calculated by subsampling to 10,000 reads per sample.

Genomic DNA (gDNA) from culturable isolates was isolated as described above. The full-length 16S gene was PCR amplified from gDNA using primers 27F and 1492R (SI Table S1). PCR reactions were run on a 0.7% agarose gel in 1xTAE, purified with NucleoSpin Gel cut and PCR clean-up kit (Macherey Nagel, Germany), and the entirety of each amplicon was sequenced using Oxford Nanopore Technologies sequencing (Oxford, UK). Annotated alkane 1-monooxygenase (*alkB*) was identified in the sequenced genome of *Rhodococcus aetherivorans* strain DMU1 (accession no. CP050952.1) and an alkene monooxygenase was identified in the sequenced genome of *Rhodococcus aetherivorans* strain 11-3 (accession no. CP080406.1). Primers were designed to amplify the entire coding region from isolated genomic DNA (SI Table S1). PCR reactions were run on a 0.7% agarose gel in 1xTAE, purified with NucleoSpin Gel cut and PCR clean-up kit (Macherey Nagel, Germany), and the entirety of each amplicon was sequenced using Oxford Nanopore Technologies sequencing (Oxford, UK). Sequences were then compared via BLASTn to identify each strain of *Rhodococcus* sp. isolates.

## Results & Discussion

### Initial enrichment strategy and collection of microbial consortia on plastic-enriched soil

Samples were collected from the yard of a local recycling plant or from cooling tank sludge and incubated with LDPE sheets in a minimal medium supplemented with 0.05% yeast extract for seven months until robust biofilms formed on the sheets. The original goal of the incubation with LDPE was to identify microbes that could degrade the intact pristine polymers. However, we did not identify degradation of the LDPE sheets using various techniques including FTIR, scanning electron microscopy, contact angle measurements, and gel permeation chromatography (GPC) (data not shown). We reasoned that biofilms that formed on the surface of LDPE sheets over a period of several months in a very low nutrient environment would have adapted to utilize linear alkanes, which are found in LDPE sheets^30^ and likely leach into the medium. We, therefore, chose to deconstruct the recalcitrant PE polymer using hydrogenolysis over Ru/C catalyst and then utilize the biofilms formed on the LDPE sheets to valorize the hydrogenolysis products via enzymatic functionalization. A diagram of the sampling, collection and assay methods is shown in Figure 2. The biofilms were collected in a low-salt solution and stored for future use. Seven consortia were then tested for alkane degradation capabilities, using *n*-hexadecane as a model compound (data not shown). Two consortia, hereafter referred to as parent 1 (P1) and parent 2 (P2), that grew well on *n*-hexadecane were selected for further analysis. The selected consortia were utilized in two ways: 1) in an enrichment strategy to enrich for increased catabolism of linear alkanes and 2) in alkane degradation assays of a PE-deconstruction product mix produced via hydrogenolysis of HDPE. DMSO stocks of consortia 1 and 2 were used as inocula for the enrichment strategy to enrich for alkane-catabolizing species. The consortia were grown for one week in Bushnell-Haas medium with a commercially-available linear alkane mix of even-numbered alkanes C_10_-C_40_ as the sole carbon source. The consortia were subjected to two rounds of passaging in the same growth conditions for two and four weeks. The entire culture of each enriched consortium 1 and 2, hereafter referred to as enriched 1 (E1) and enriched 2 (E2), was collected and stored as DMSO stocks for future use. The original (parent) consortia P1 and P2 and the enriched consortia E1 and E2 were then grown on the PE-deconstruction product mix to determine the fractional degradation and conversion of the alkanes. Additionally, the genus-level bacterial community structure of parent and enriched consortia before and after growth on the PE-deconstruction product mix were obtained via 16S sequencing. We then identified and quantified the metabolites derived from *n*-hexadecane as a model alkane.

**Figure 2.**
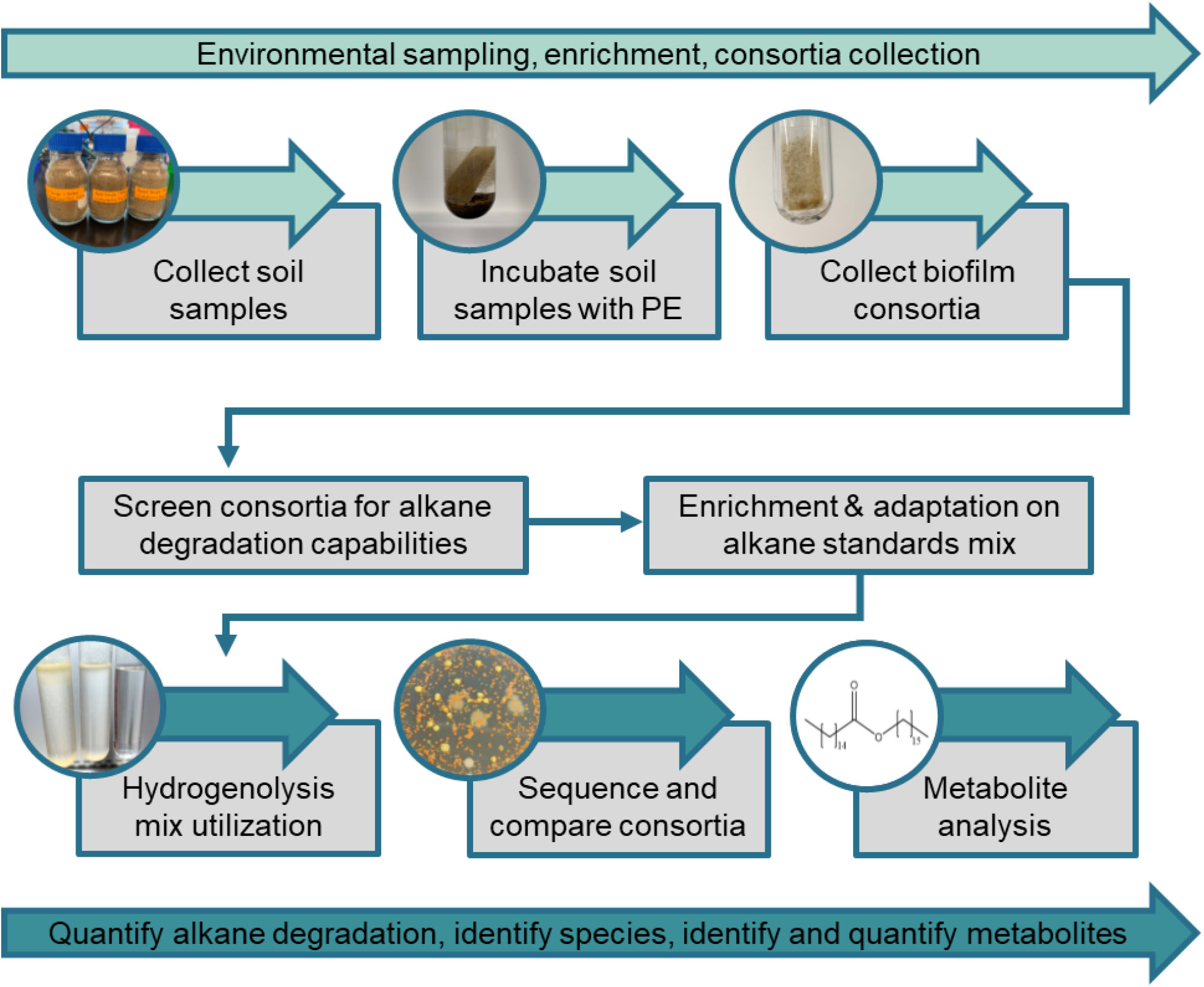
Sampling, Enrichment and Growth schematic. Environmental soil samples were collected from the yard of a local recycling plant and incubated with low density polyethylene (LDPE) sheets in minimal medium. Biofilms that formed on the plastic sheets were collected in 0.9% NaCl and stored at −80°C in 7% DMSO or 20% glycerol. Consortia were tested for growth on a model linear alkane, *n*-hexadecane. Two consortia, 1 and 2, were selected for further testing. The consortia then underwent an enrichment and adaptation procedure consisting of serial passaging with a commercial alkane standards mix as the sole carbon source for a total of 7 weeks. The parent consortia and the evolved consortia were then grown on the PE deconstruction product mix as the sole carbon source and alkane degradation and conversion were compared. 16S sequencing was performed on the consortia before and after enrichment to identify the species in each consortium. The metabolites produced by each evolved consortium were identified and then quantified after growth on model alkane hexadecane.

### Quantification of PE hydrogenolysis alkane product degradation after microbial treatment

After a subculturing procedure to increase the biomass of each consortium (Figure 3), P1 and P2 were grown with the PE-deconstruction product mix for 14 days. The remaining alkanes were then extracted two times from the entire culture with 5 mL of DCM total to quantify the fractional degradation of each chain length. Extractions from negative controls without bacterial inocula were performed to quantify alkanes lost during the incubation and extraction procedures. A sample of the pure input PE-deconstruction product mix was also prepared at the time of the extraction for comparison. The alkane concentrations present in the pure input PE-deconstruction product mix, and the residual alkane concentrations present in DCM extractions of the incubated negative controls and consortia cultures (Figure 4A) show, as expected, that volatile, small-chain alkanes are absent in the negative control and consortia-treated cultures, presumably due to evaporation. A fraction of the alkanes is also lost during the incubation and the extraction procedure itself, as seen when comparing the pure mix to the negative controls (Figure 4A). The conversion by chain length and the total were then calculated to account for the loss in the negative control (see Methods).

**Figure 3.**
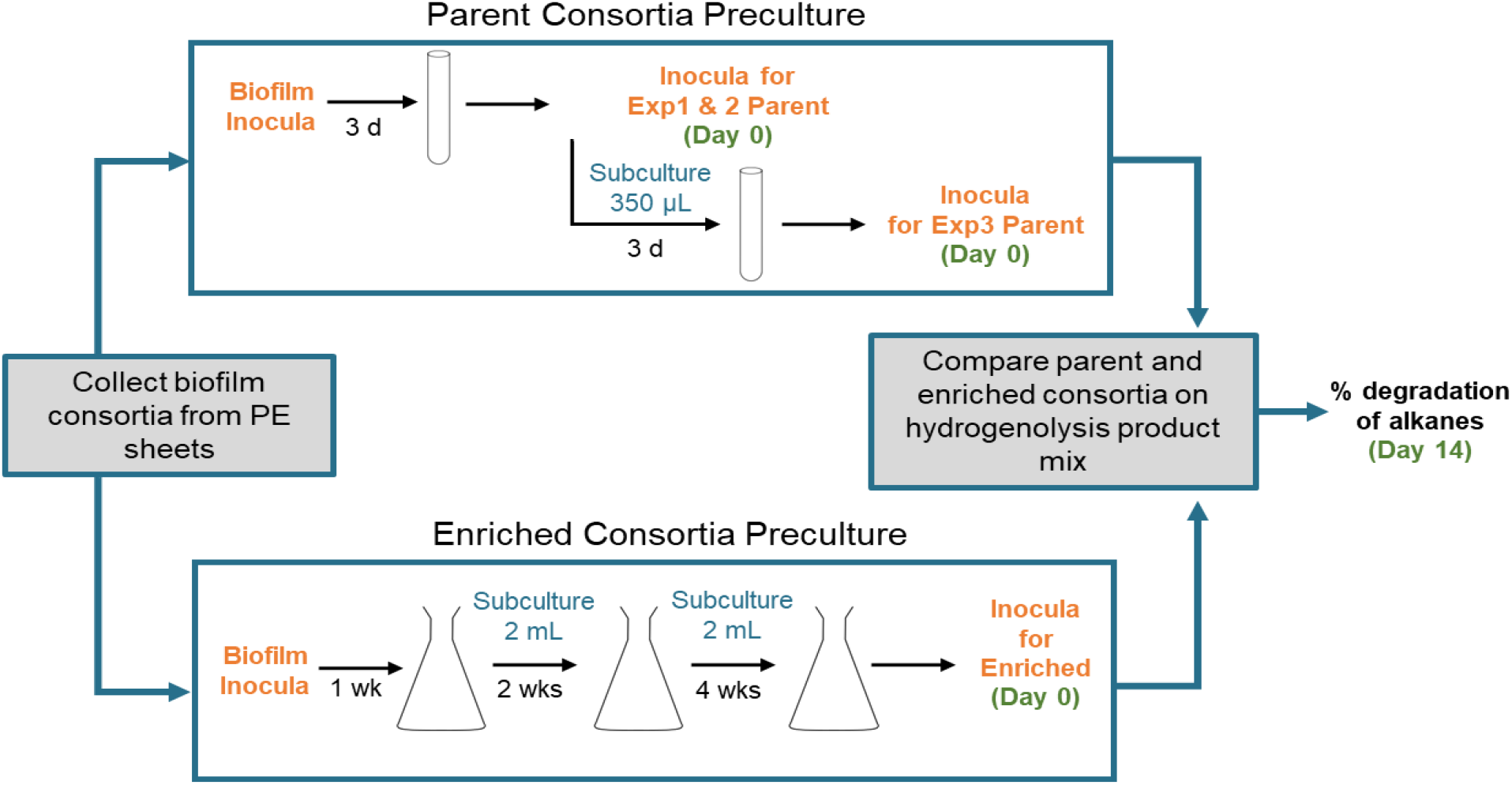
Preculture procedures for parent and enriched consortia. Biofilms collected from PE sheets were stored at −80°C in 7% DMSO or 20% glycerol. Biomass was increased using a preculture procedure for the parent consortia. The biofilm glycerol stocks were used to inoculate Bushnell-Haas (BH) medium with the PE hydrogenolysis product mix as the sole carbon source. Precultures for parent consortia Exp1 and Exp2 were incubated for 3 days and then stored as 7% DMSO stocks for future use. The preculture for Exp1 was subcultured in BH medium with the PE deconstruction product mix as the sole carbon source for 3 days, then stored as 7% DMSO stocks for future use (Exp3). The biofilm collected from PE sheets was also enriched using a Restek alkane standards mix as the sole carbon source. Consortia 1 and 2 were cultured for one week in BH medium with the Restek mix as the sole carbon source. The consortia were then subcultured twice, for two and four weeks, in the same medium with Restek mix for a total of 7 weeks enrichment on linear alkanes. Using DMSO stocks made after the preculture procedure, the parent consortia and the evolved consortia were then grown on the PE deconstruction product mix as the sole carbon source and alkane degradation and conversion were compared. 16S sequencing was performed on the consortia before (Day 0) and after (Day 14) enrichment to identify the species in each consortium. Stocks of the consortia, which were used as inocula in subsequent steps, are indicated in orange. Subculture volumes are indicated in blue. Samples that were utilized for 16S sequencing (Day 0 and Day 14) are indicated in green.

**Figure 4.**
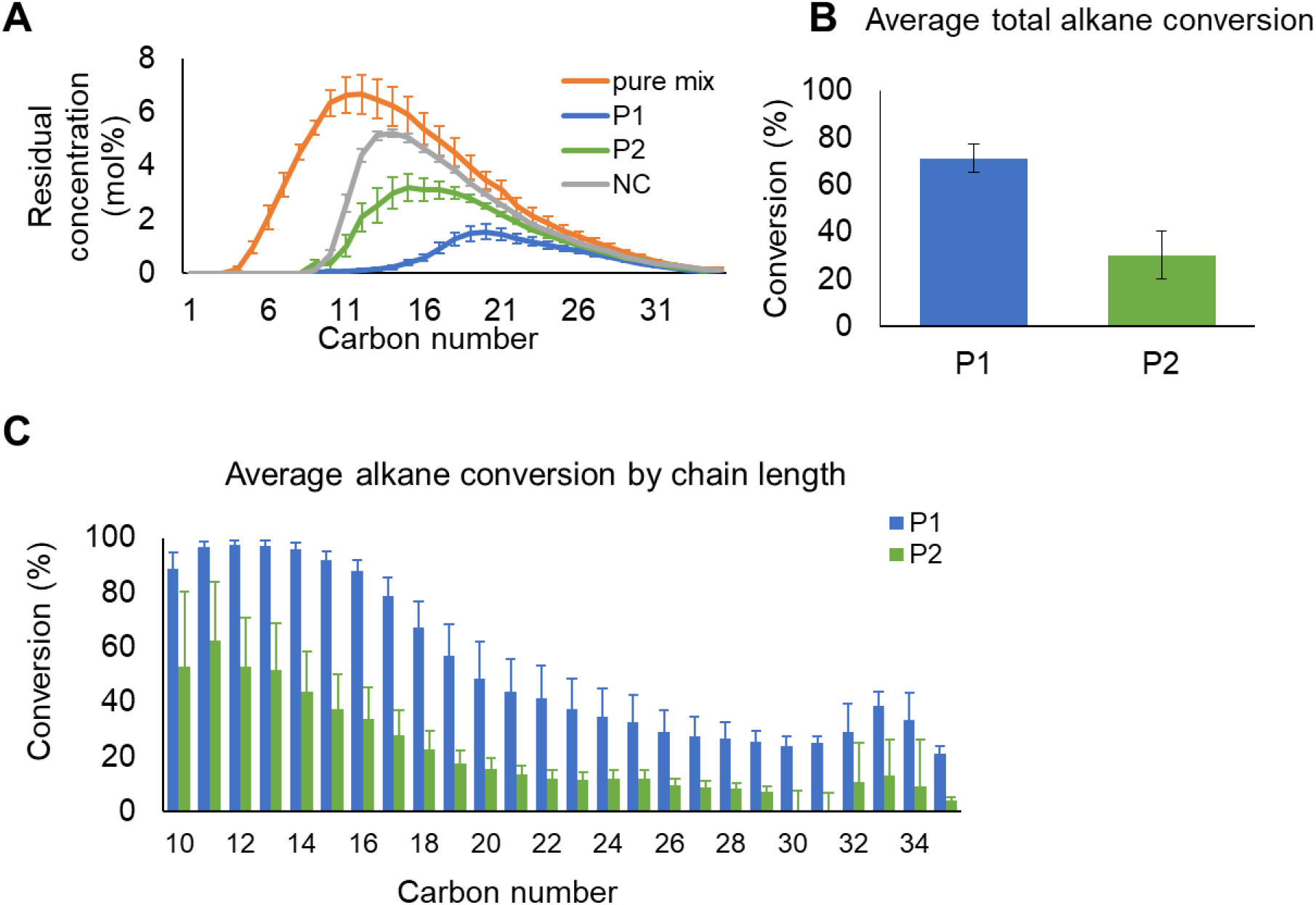
(A) Residual concentrations of linear alkanes after incubation with parent microbial consortia P1 or P2 and an untreated negative control (NC), along with the liquid product distributions for the hydrogenolysis reaction of HDPE (MW= 76 kDa) over a Ru/C catalyst. Quantification of PE deconstruction product mix degradation after microbial treatment by P1 or P2 for two experiments with two independent cultures each shown as (B) total carbon conversion or (C) carbon conversion of individual alkane chain lengths. Error bars indicate standard error of the mean (SEM).

The average total conversion by P1 was 71% and the average total conversion by P2 was 30% (Figure 4B). Both consortia have higher conversions of chain length < C_20_ than larger chain length alkanes, with P1 performing better than P2 across all chain lengths (Figure 4C). Conversion of chain lengths < C_20_ by P1 ranged from to 98% of C_13_ to 56% conversion of C_19_. P1 gave a conversion of ranging from 48% of C_20_ to 22% conversion of C_35_ (Figure 4C). The highest conversion by P2 was 61% of C_13_ (Figure 4C). In P2, the conversion was less than 20% for chain lengths of C_20_ or longer, and less than 10% for chain lengths of C_28_ or longer (Figure 4C). Although we used the same preculture procedure for experiments 1 and 2, we saw a stark difference in alkane conversion by P2 cultures in Exp2 as compared to Exp1 (Figure S1A & B), with average total alkane conversion dropping to 13% (Figure S1). We then did a subculture of Exp1 as the preculture (see Figure 3). We found that subculturing produced results similar to the original preculture procedure, with an average total conversion of alkanes by P2 of 47% in Exp1 and 42% in Exp3. Conversion of each alkane chain length was also similar between Exp1 and Exp3 (see Figures S1A & C). The preculture method to increase biomass likely resulted in differential species enrichment in the community structures of the cultures of Exp1 and Exp2, and therefore differences in alkane conversion.

### Quantification of the conversion of PE-hydrogenolysis alkane products using microbial consortia enriched on standard alkane mix

To simplify the consortia by reducing the number of species and enrich for species with increased alkane utilization via enrichment on linear alkanes, we performed serial batch passaging of each consortium on minimal medium with a commercially-available mix of alkane standards, containing even-numbered n-alkanes (C_10_-C_40_), as the sole carbon source (see Figures 2 & 3). Consortia were grown for one week on the alkane standards mix and then subcultured into fresh medium. This process was repeated after two weeks and then four weeks of growth on the alkane standards mix (Figure 3). The enriched consortia, hereafter referred to as E1 or E2, were grown on the PE-deconstruction product mix and compared to the parent consortia (Figure 5). The residual concentration of alkanes extracted from each culture, along with the input PE-deconstruction product alkanes and the negative control (Figure 5A), show that the total alkane conversion by E1 increased from 71% to 85% (Figure 5B). E2 had a significantly increased total alkane conversion from 30% to 86% (Figure 5B). E1 performed better than the parental consortium on all alkane chain lengths (Figure 5C), with an average 2-fold increase in the utilization of alkanes with C_20_ chain lengths or longer (Figure 5C). E2 had marked improvements in alkane utilization, with 4- to 12-fold increase in the utilization of alkanes with C_20_ chain lengths or larger (Figure 5D). The alkane enrichment procedure likely enriched for microbes with greater alkane utilization capabilities, as evidenced by the increase in total alkane utilization by both consortia (Figure 5B). Note that E2 had marked increases in utilization of *all chain lengths*, likely due to a simplified consortium and enrichment in species that can utilize longer-chain alkanes. We discuss below the microbial community structure of each consortium and relate this to alkane utilization profiles.

**Figure 5.**
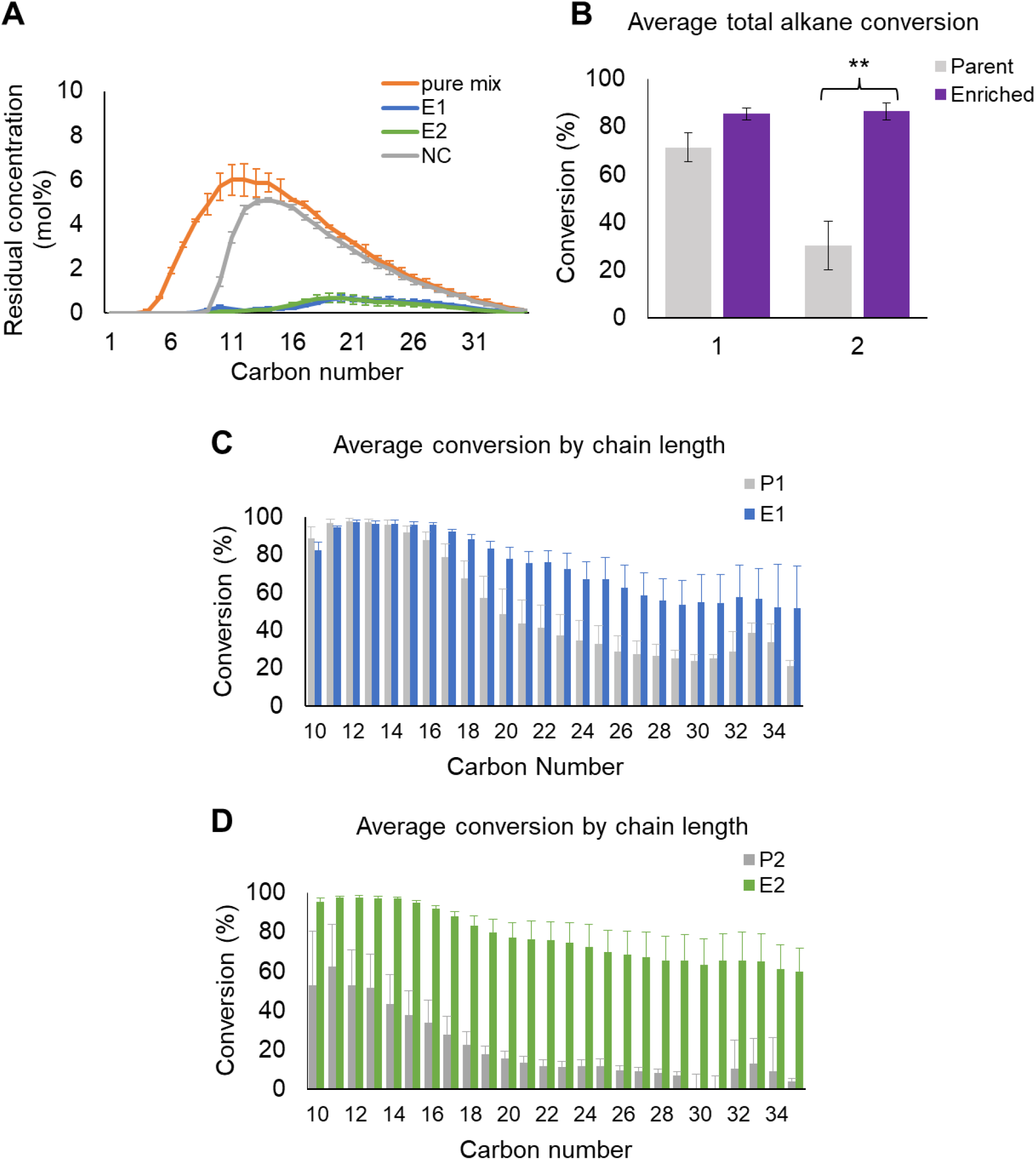
(A) Quantification of PE deconstruction product mix degradation after microbial treatment with enriched consortium 1 (E1) or consortium 2 (E2) for 14 days. (B) Comparison of average total carbon conversion by parent and alkane-enriched consortia E1 and E2. Comparison of average carbon conversion of individual alkane chain lengths by parent and alkane-enriched consortia (C) E1 or (D) E2. All figures display the average of two independent experiments with two independent cultures each with SEM indicated by error bars. Statistics were calculated using an unpaired Student’s t-test; **, p<0.005.

### Metabolite analysis after growth on model alkane hexadecane

Bacterial utilization of the PE-hydrogenolysis alkanes under normal growth conditions likely results in the production of biomass and CO_2_, as fatty acids produced from linear alkanes are funneled into the β-oxidation pathway^37, 38^. Metabolic pathways for aerobic linear alkane degradation have been well-characterized^14, 16, 37–39^. The first step in the pathway is catalyzed by alkane hydroxylases, which perform the initial oxidation of the n-alkane. There are several unrelated classes of alkane hydroxylases that have been characterized^37–40^. The typical range of substrates for each class of alkane hydroxylase is as follows: methane monooxygenases, C_1_-C_4_ alkanes^41^; alkane monooxygenases (AlkB-like monooxygenases), C_5_-C_32_^16, 39^; bacterial P450s, C_5_-C_16_ alkanes^42, 43^; eukaryotic P450s, C_10_-C_16_^44^; and dioxygenases, C_10_-C_30_ alkanes^45^. These enzymes are either rubredoxin-dependent or cytochrome P450 monooxygenases. Other enzymes with activity on longer chain-length alkanes include LadA, a flavoprotein-dependent monooxygenase with activity on C_15_-C_36_^46, 47^, and AlmA, a flavin-binding monooxygenase with activity on C_20_-C_32_+ alkanes^48^. Homologs of AlmA have been found in several species^37^. Degradation of n-alkanes up to chain length C_44_ has been reported^49, 50^.

There are four routes for the initial oxidation that differ based on the position of the carbon atom that is attacked by the enzyme: terminal, biterminal, subterminal and the Finnerty pathway^51–55^. Each pathway culminates in the formation of a carboxylic acid which enters β-oxidation to yield acetyl-CoA^51^. However, under nitrogen limitation, many bacterial species will convert alkanes into carbon storage molecules such as wax esters, triacylglycerols (TAGs), and polyhydroxyalkanoates (PHAs)^37, 38, 56^. Wax esters, neutral lipids formed from the esterification of a long-chain acyl-CoA molecule with a long-chain alcohol (Figure 6), are widely used in pharmaceuticals, cosmetics, lubricants, and food industries^37^. We first identified the metabolites produced by E1 without nitrogen limitation to establish that each consortium was oxidizing the n-alkane substrate and funneling this into the β-oxidation pathway. We used *n*-hexadecane as a model substrate on which to test metabolite production.

**Figure 6.**
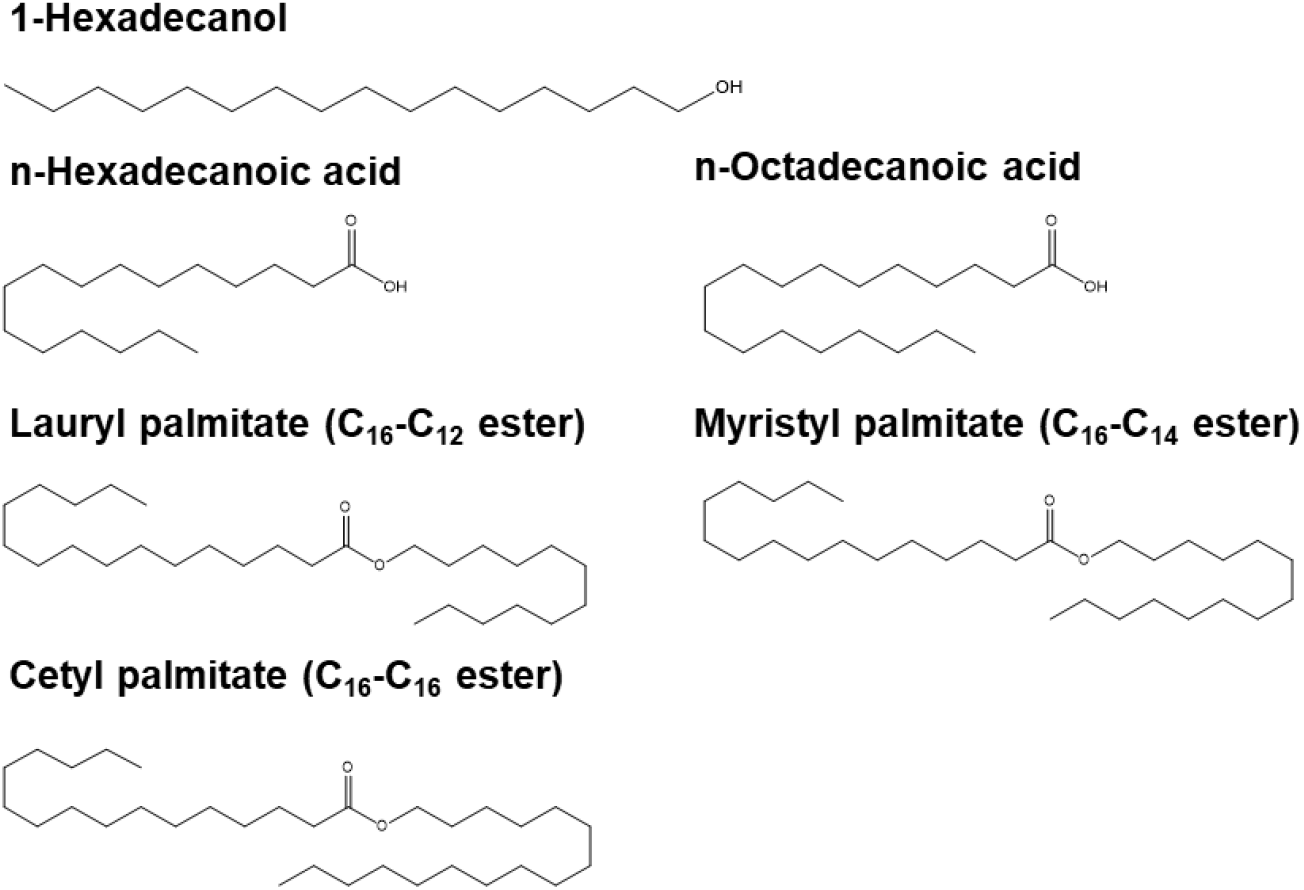
Structures of compounds identified via GC-MS of organic phase extracted from nitrogen-limited cultures of E1 and E2.

We found that after 24 hours of E1 and E2 incubation with hexadecane, hexadecanol was formed, and after 48 hours, hexadecanoic acid was formed (data not shown); that is, terminal oxidation occurs. From the perspective of plastics recycling and valorization, the biological production of lipid storage molecules from PE hydrogenolysis products is a good way to preserve the longer alkane chain lengths and to minimize the production of CO_2_ by preventing the complete breakdown of alkanes via β-oxidation. Using *n*-hexadecane, we determined that both E1 and E2 both catalyzed the formation of wax esters after 6 and 14 days of incubation (data not shown). We then quantified wax ester production and the production of additional metabolites after 14 days of incubation in a nitrogen-limited medium (Table 1, Figure S2). Cetyl palmitate (C_16_-C_16_ ester) was the highest-produced metabolite by E1 (35.6 mg/gCDW) and E2 (30.18 mg/gCDW) (Table 1), and 1-hexadecanol was the second highest-produced metabolite (Table 1). Hexadecanoic and octadecanoic acid, lauryl palmitate (C_16_-C_12_ ester) and myristyl palmitate (C_16_-C_14_ ester) (Table 1) were also identified (see Figure 6).

**Table 1.** Metabolites identified in nitrogen-limited cultures of E1 and E2

|  | **E1 (mg/gCDW)** | **E2 (mg/gCDW)** |
| --- | --- | --- |
| 1-Hexadecanol | 9.745 ± 3.417 | 13.849 ± 3.062 |
| Hexadecanoic acid | 2.420 ± 0.185 | 2.527 ± 0.239 |
| Octadecanoic acid | 2.060 ± 0.285 | 1.686 ± 0.255 |
| Lauryl palmitate | 0.684 ± 0.093 | 1.787 ± 0.197 |
| Myristyl palmitate | 3.600 ± 0.492 | 6.006 ± 0.656 |
| Cetyl palmitate | 35.578 ± 2.910 | 30.175 ± 3.027 |

### Microbial diversity of environmental consortia

To understand the differences in alkane utilization by the parent and enriched consortia, we performed 16S v4 sequencing to characterize the bacterial community structure at the genus level. Sequenced samples from the consortia had between 23 and 61 identified bacterial genera (SI Tables S2 & S3). To simplify the analyses, we selected genera representing at least 3% relative abundance in any sample. Genera not meeting this threshold were aggregated into a single “Other genera” category. This threshold resulted in 24 genera that were compared among the consortia, but not every genus was represented in every sample sequenced. Five of these were unclassified genera from the families *Phyllobacteriaceae*, *Bacillaceae*, *Planococcaceae*, *Alphaproteobacteria* and *Gemmatimonadaceae*.

As expected, the microbial community structure of each experiment of the parent consortia varied due to the use of a different preculture for each (Figure S3). The most striking difference in community composition across the three P1 experiments is the relative abundances of *Hydrogenophaga*, *Sphingopyxis* and *Bacillus*, particularly when compared to the community composition at day 14 after growth on the PE-deconstruction product mix. P1 Exp1 and Exp3 both have higher relative abundances of *Bacillus* than is seen in any other sample. P1 Exp2 is dominated by *Hydrogenophaga*, which makes up over 70% of the community at day 0. By day 14, the ratios of *Hydrogenophaga* and *Sphingopyxis* are all more similar across all P1 experiments (Figure S3). P1 Exp1 and Exp3 had similar conversion of total alkanes and of individual chain lengths (Figure S1). P1 Exp2 performed similarly to other experiments on shorter chain alkanes, but not as well on medium and long chain alkanes (Figure S1). The differences in alkane conversion among the P1 experiments indicate that starting species ratios are important. It is also possible that the higher relative abundance of *Bacillus* in Exp1 of P1 played a role in better total alkane conversion (Figure S1). However, negligible numbers of *Bacillus* were present in the enriched consortia at both days 0 and 14. This does not rule out *Bacillus* as an important genus for alkane conversion in P1 Exp1. Differences in microbial consortia among P1 experiments were likely caused by a variety of factors during the growth and enrichment procedures, including competition among species. We noted that P2 Exp2, which had the lowest alkane conversion rate of any of the parent trials (Figure S1), had a lower relative abundance of *Sphingopyxis* (6%) at day 14 than the other experiments of both P1 (26%, 10%, 13%) and P2 (39% and 19%) (Figure S3). P2 Exp2 starts with very low relative abundances of both *Sphingpyxis* and *Hydrogenophaga*, which may explain why the alkanes with chain lengths > C20 are not converted as well as in Exp2 (Figure S1). *Bordetella* has high relative abundances in each P2 experiment after 14 days, while we see relatively less *Bordetella* in P1 (Figure S3). An unclassified *Bacillaceae* genus also dominates the communities of each P2 experiment at day 0. However, as seen in P1 with the *Bacillus* genus, the unclassified *Bacillaceae* genus is present in negligible numbers. This example could also explain why P2 did not perform as well as P1 in the alkane conversion experiments. The endpoint samples of P1 and P2 are similar, but starting community compositions vary greatly. P2 experiments have lower numbers of important alkane degrading genera at day 0 (Figure S3), and therefore P2 did not convert the alkanes as well as P1 (Figure S1). As with consortium P1, day 14 samples of P2 are surprisingly similar to each other, given the differences in community composition at the start of the experiment. Given time, the species ratios may have evened out, and resulted in the same level of conversion seen in the best-performing experiments. The differences in alkane conversion seen between the three experiments of P1 and P2 (Figure S1) indicate that optimal starting ratios are important for conversion of all alkane chain lengths.

The enrichment procedure on the alkane standards mix enriched for *Hydrogenophaga* and *Rhodococcus* genera in both consortia, as seen by the high relative abundances of each genus at day 0 in both E1 and E2 (Figure 7). After growth on the PE-deconstruction product mix for 14 days, *Sphingopyxis* and *Phyllobacteriaceae* are enriched in both consortia, *Pseudoxanthomonas* and *Parvibaculum* are enriched in E1, while *Devosia* is enriched in E2. The relative abundance of *Hydrogenophaga* decreases from day 0 to day 14. The relative abundances of *Hydrogenophaga* in all endpoint sample of both parent and enriched consortia are similar, indicating that there may be an optimal ratio of *Hydrogenophaga* to other consortium members.

**Figure 7.**
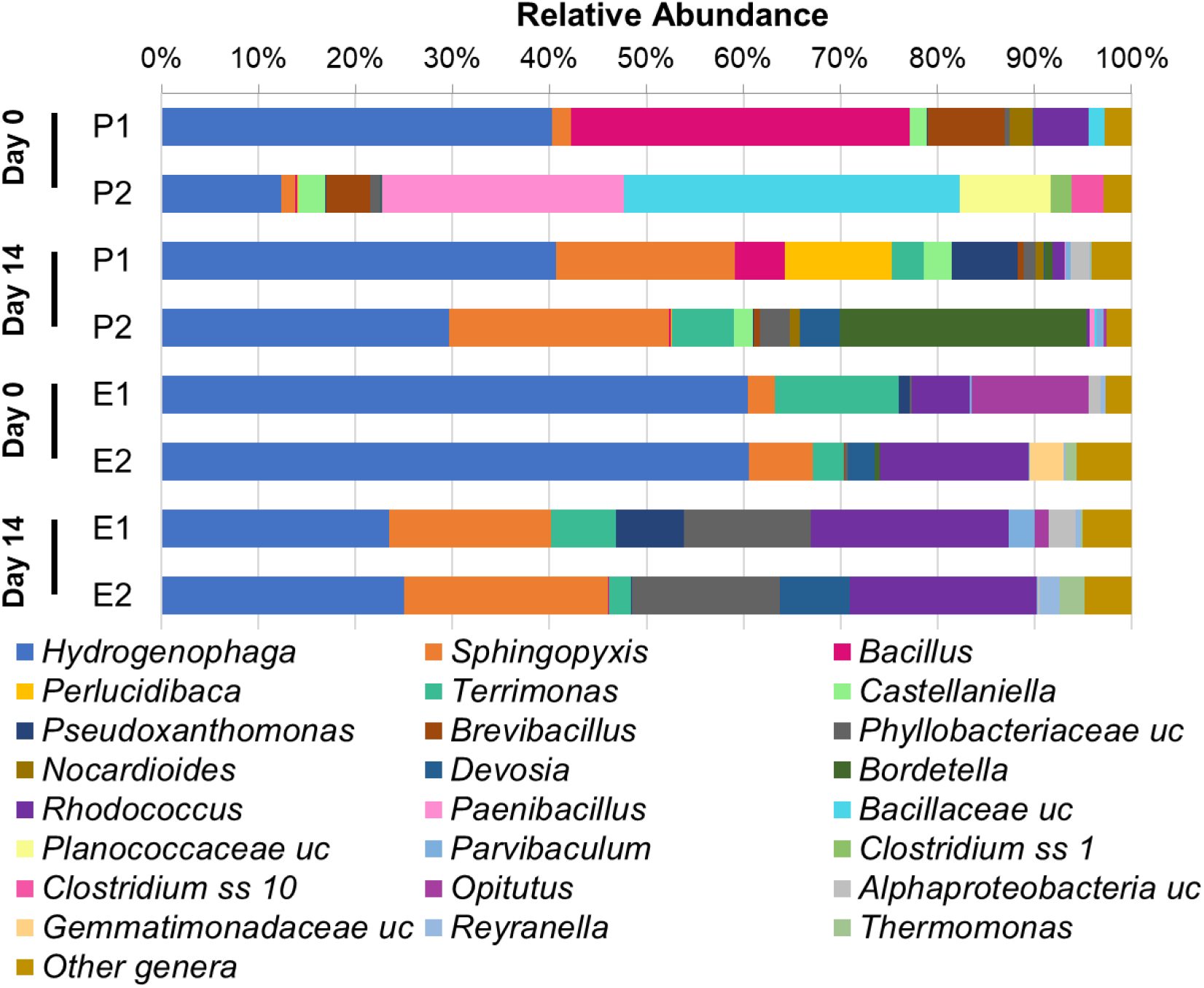
(A) Genus-level bacterial community structures identified in Parent consortia 1 and 2 (P1, P2) and enriched consortia 1 and 2 (E1, E2) at initial inoculum (day 0) and after incubation with PE deconstruction product mix for 14 days. Relative abundances are expressed as a percentage of DNA sequence reads. uc = unclassified genera, ss = sensu stricto.

*Hydrogenophaga* and *Sphingopyxis* are two of the most important genera for degradation; together they make up more than 50% of the relative abundance of each consortium on day 14 for both parent and enriched consortia (Figure 7). Other important genera include *Rhodococcus* and an unclassified *Phyllobacteriaceae* genus, whose relative abundances increased in the enriched consortia samples (Figure 7). Members of the *Hydrogenophaga* genus, formerly classified as pseudomonads, are yellow-pigmented Gram-negative bacteria that consume hydrogen^57^. *Hydrogenophaga* species also degrade linear, branched, and cyclic alkanes and polycyclic aromatic hydrocarbons (PAHs) and have been identified in environmental samples from hydrocarbon-contaminated sites^58–62^. *Sphingopyxis* is enriched in all endpoint samples for both parent and enriched consortia (Figure 7, S3). Members of the *Sphingopyxis* genus are Gram-negative bacteria that are of interest for their bioremediation potential due to the wide variety of catabolism genes identified^63–65^. *Sphingopyxis* is also associated with PAH degradation^63^. Its role in linear alkane degradation has not been widely reported^66^. Accumulation of PHAs, but not wax esters, has been reported previously for *Hydrogenophaga* and *Sphingopyxis* species^67–71^. The *Rhodococcus* genus is comprised of Gram-positive bacteria that have broad metabolic versatility and are studied for bioremediation potential^72^. *Rhodococcus* species are well-known for degradation of linear, branched and cyclic alkane substrates^14, 37, 73–76^. Accumulation of wax esters and other lipid storage compounds has also been investigated in rhodococci, and notably for *Rhodococcus jostii* and *Rhodococcus opacus*^77, 78^. A recent study characterizing hydrocarbon-degrading organisms collected from sediment at the site of an oil spill found that both *Rhodococcus* and *Hydrogenophaga* were enriched in all samples tested^62^.

### Identification and investigation of isolates after enrichment on long-chain alkane tetracosane (C_24_)

Since the conversion of alkane chain lengths of > C_20_ was poor in the parent consortia, we wanted to identify isolates from each consortium that could catabolize a model long-chain alkane, tetracosane (C_24_). To this end, we first grew each parent consortium on tetracosane as the sole carbon source for 14 days to enrich for species that could utilize longer chain alkanes. We then performed dilution plating and identified 8 morphologically distinct colonies total from both consortia. Whole-gene 16S rDNA sequencing was used to identify each isolate and then isolates were tested for C_24_ catabolism capabilities (Figure S4). The isolates identified included two *Rhodococcus* sp., *Brevibacillus reuszeri*, *Bacillus thuringiensis*, *Pseudoxanthomonas indica*, *Agromyces indicus*, *Bordetella petrii*, and *Lysinibacillus macroides*. Only two isolates were capable of growth on C_24_ (Figure S4), S1 isolated from P1 and S5 isolated from P2. Both were identified as *Rhodococcus sp.* that shared 100% nucleotide identity with multiple strains, including *Rhodococcus aetherivorans* IcdP1, *Rhodococcus* sp. strain 11-3, *Rhodococcus* sp. strain DMU1, *Rhodococcus aetherivorans* N1, *Rhodococcus* sp. strain D-50, and *Rhodococcus* sp. WB1. The two *Rhodococcus* species had identical 16S sequences, but differing morphology when plated on LB agar.

To identify each isolate to the species/strain level, we sequenced additional genes amplified from each isolate. We searched the sequenced genomes of *Rhodococcus* sp. strain DMU1 and *Rhodococcus* sp. strain 11-3 for annotated alkane hydroxylases. The *alkB* alkane 1-monooxygenase gene from *Rhodococcus* sp. strain DMU1 and an alkene monooxygenase gene from *Rhodococcus* sp. strain 11-3 were selected, primers were designed to amplify the genes from the genomic DNA of each *Rhodococcus* sp. isolate (SI Table S1), and whole-gene sequencing was performed as for the 16S rRNA gene. The sequenced *alkB* gene from S1 shared 99.92% nucleotide identity with DMU1 and 99.84% nucleotide identity with *R. aetherivorans* IcdP1. The sequenced alkene monooxygenase gene from S1 shared 100% nucleotide identity with *R. aetherivorans* IcdP1. Given the results of the whole-gene sequencing of these monooxygenases, it is likely that S1 is *R. aetherivorans* IcdP1. The sequenced alkane monooxygenase gene from S5 shared 99.5% identity with *R. aetherivorans* N1 and 99.42% identity with *Rhodococcus* sp. strain 11-3. The sequenced alkene monooxygenase shared 99.66% identity with *R. aetherivorans* N1 and 99.58% identity with *Rhodococcus* sp. strain 11-3. S5 is likely *R. aetherivorans* N1, but additional sequencing is required to confirm this finding. Based on these sequences, we can conclude that isolate S1 and isolate S5 are different strains, likely both *Rhodococcus aetherivorans*. It is of interest to note that another *R. aetherivorans* strain, *R. aetherivorans* BCP1, has been reported to grow on *n*-alkanes up to C_28_^79^. However, promoter studies indicated that the *alkB* gene in this strain is not induced by alkanes larger than C_22_^79^. The *alkB* genes from *R. aetherivorans* IcdP1 and *R. aetherivorans* N1 share 99.59% nucleotide identity with the *alkB* gene in *R. aetherivorans* BCP1. The *alkB* gene found in *R. aetherivorans* IcdP1 is disrupted due to a premature stop codon. The produced enzyme may still be active on alkanes, as only the last 16 amino acids of the C terminus are missing. Given that the strains we isolated utilize both C_24_ and C_36_ n-alkanes, it is likely that additional alkane hydroxylases that are induced by and can functionalize larger alkanes are present in the genome of *R. aetherivorans* strains IcdP1 and N1 and further investigation and enzyme characterization is required.

The bacterial community structure analyses revealed that *Rhodococcus* was enriched in both E1 and E2, while *Pseudoxanthomonas* was enriched in E1 as compared to the parent consortia (Figure 7). The data suggest that these genera may be important for long-chain alkane utilization, as utilization was increased after the enrichment procedure (Figure 5). Interestingly, although a *Pseudoxanthomonas* isolate was identified after additional consortia enrichment on C_24_, the isolate was not capable of growth on C_24_ as the sole carbon source. This suggests that the *Rhodococcus aetherivorans* isolates, the only two isolates capable of growth on C_24_ as the sole carbon source, are important for the initial functionalization of long-chain alkanes, while other genera in the consortia likely utilize intermediates produced during alkane degradation. Only morphologically distinct, aerobic culturable isolates were tested for long-chain alkane degradation capabilities. Other species found in the consortia likely also degrade long-chain alkanes, including those identified in the community structure profiling such as *Hydrogenophaga* and *Sphingopyxis*. While other *Rhodococcus* spp. have been reported to produce wax esters^77, 78^, the accumulation of wax esters by *Rhodococcus aetherivorans* has not been previously reported. However, *R. aetherivorans* has been shown to produce TAGs and PHAs from a toluene feedstock, highlighting the utility of this species in bioremediation and valorization of recalcitrant environmental contaminants^80, 81^.

### Implications and the path forward

Here we have demonstrated that we can tune microbial consortia using an adaptation and enrichment procedure to increase conversion of a PE deconstruction product mix. The same procedure works with other polymer deconstruction product mixes, such as branched alkanes produced via polypropylene deconstruction (data not shown). Thermodynamic differences among polymer types limit mechanical recycling, which involves heat treatment to produce secondary, raw polymer materials^82, 83^. Mixed-stream recycled plastics need to be separated and each polymer type treated individually, which is not always possible due to a variety of factors, including inefficient separation methods and multilayer plastic products^84–86^. We envision bioconversion of mixed deconstruction products from different polymers and tunable microbial consortia to treat complex feedstocks to valorize plastics waste. Defined consortia could be assembled by selecting *n*-alkane degrading bacterial species identified here and species known for degradation of branched alkanes or other deconstruction products. In this way, consortia could be tuned for specific mixes of waste plastics.

The microbial consortia used here performed selective terminal oxidation of linear alkanes. This is in contrast to recently published methods, such as non-thermal plasma functionalization of alkanes^87^, which does not allow for the same selectivity. Alkane hydroxylases also perform sub-terminal and bi-terminal oxidation of alkanes^51–54^. Enriching additional consortia that perform sub-terminal or bi-terminal functionalization of alkanes will allow us to tune the bioconversion step for desired end-products. Natural or engineered alkane hydroxylases are reported to produce α, ω-diols or diacids up to chain length C_16_^88–90^. Polycondensation reactions with long-chain diols and diacids produce polyesters with more-desirable properties than those produced by short- and medium-chain difunctional molecules^91^. These properties include higher polymer melting points, higher crystallinity, and increased stability against hydrolytic degradation^91^. The polycondensates’ properties become polyethylene-like as the length of the monomer chains increases^92–94^. We have identified two strains of *R. aetherivorans* that terminally oxidize long-chain alkanes. Preliminary bioinformatics analyses indicate that AlkB-like monooxygenases and cytochrome P450 monooxygenases are encoded in the genomes of both strains. These enzymes can be characterized and engineered to produce long-chain diols and diacids for production of polyethylene-like polyesters.

## Acknowledgments

This work was supported as part of the Center for Plastics Innovation, an Energy Frontier Research Center funded by the US Dept. of Energy, Office of Science, Office of Basic Energy Sciences, under award number DE-SC0021166.

