## Supplemental info for "Polyethylene valorization by combined chemical catalysis with bioconversion by plastic-enriched microbial consortia"

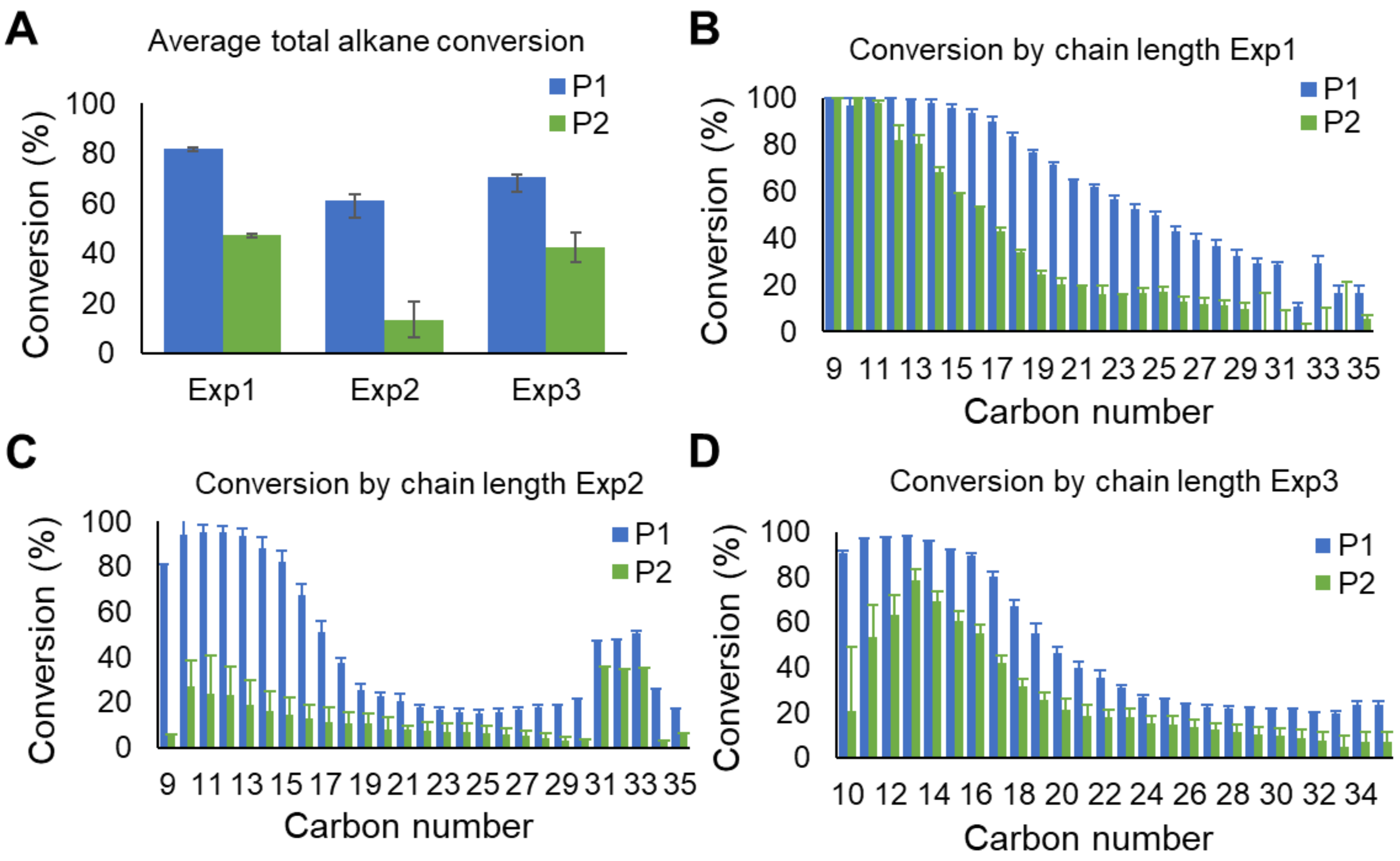


**Figure S1.** (A) Total carbon conversion or (B-D) carbon conversion of individual alkane chain lengths by P1 or P2 for three experiments with two independent culture replicates each. Error bars indicate standard error of the mean (SEM).

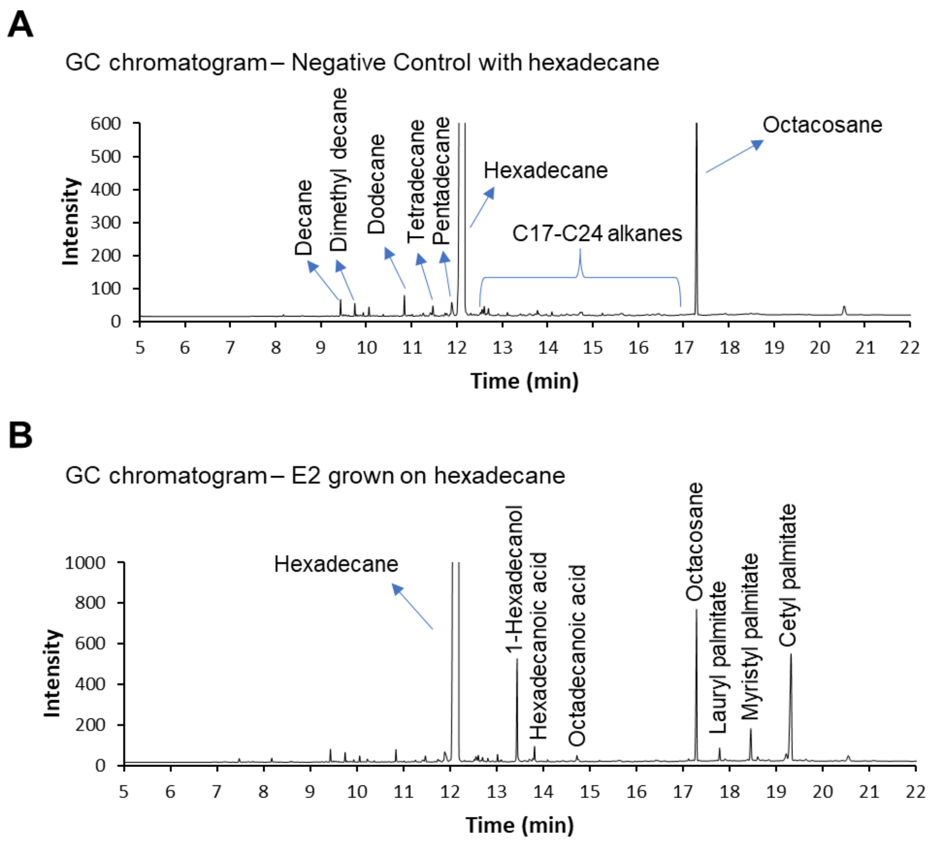


**Figure S2.** Representative GC chromatograms with peak assignments based on GC-MS identification of (A) a negative control culture with no inoculum or (B) metabolites produced by E2 when grown on hexadecane as the sole carbon source under nitrogen limitation.

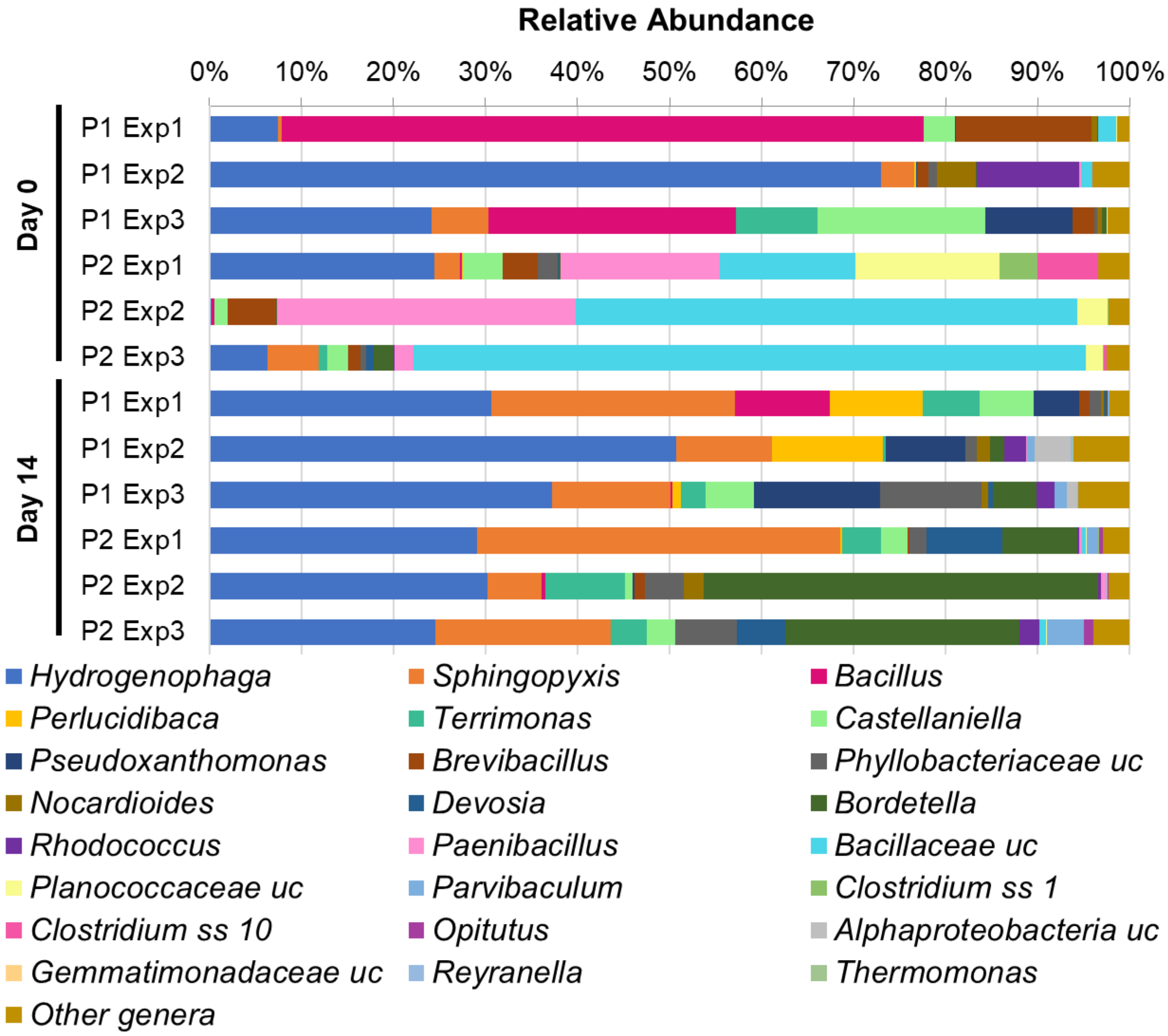


**Figure S3.** Relative abundance of bacterial genera in samples from independent Parent consortia 1 and 2 (P1, P2) experiments at initial inoculum (day 0) and after incubation with PE deconstruction product mix for 14 days. Relative abundances are expressed as a percentage of DNA sequence reads.

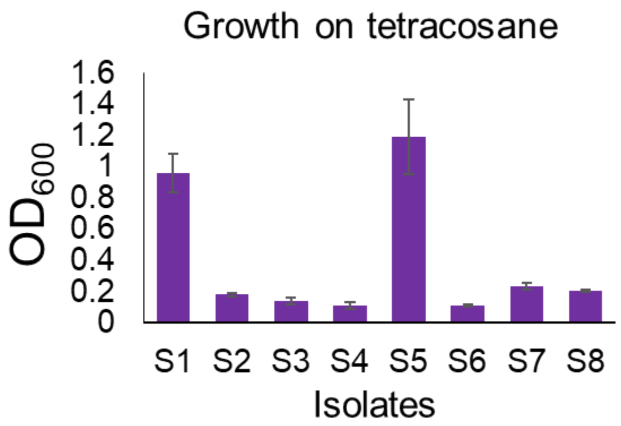


**Figure S4.** Final $OD_{600}$ of each of eight isolates grown on tetracosane for one week.

**SI Table S1.** Primers used in this study

| Primer | Sequence (5’-3’) | Length (bp) |
|---|---|---|
| 16S 27F | AGAGTTTGATCCTGGCTCAG | 1465 |
| 16S 1492R | GGTTACCTTGTTACGACTT | |
| DMU1 AMO F | CGCGTTGCAGACATATGGGA | 1260 |
| DMU1 AMO R | GATTCGACTGCTCCACCTGG | |
| 11-3 AMO F | TACGACAAGTTCACCAGCCC | 1236 |
| 11-3 AMO R | ACGCCACACATGTTGGAGAA | |

**SI Table S2.** Relative genera abundance and number of genera identified in samples from each experiment (1, 2 and 3) of consortia P1 and P2 at Day 0 and Day 14.

| | **Day 0** | | | | | | **Day 14** | | | | | |
|---|---|---|---|---|---|---|---|---|---|---|---|---|
| **sample** | **P1 Exp1** | **P1 Exp2** | **P1 Exp3** | **P2 Exp1** | **P2 Exp2** | **P2 Exp3** | **P1 Exp1** | **P1 Exp2** | **P1 Exp3** | **P2 Exp1** | **P2 Exp2** | **P2 Exp3** |
| *Acetobacteraceae* unclassified | 0.00 | 0.02 | 0.00 | 0.00 | 0.00 | 0.00 | 0.00 | 0.00 | 0.01 | 0.00 | 0.00 | 0.00 |
| *Acidimicrobiales* unclassified | 0.00 | 0.01 | 0.00 | 0.00 | 0.00 | 0.00 | 0.00 | 0.00 | 0.00 | 0.00 | 0.00 | 0.00 |
| *Actinobacteria* unclassified | 0.00 | 0.01 | 0.00 | 0.00 | 0.00 | 0.00 | 0.00 | 0.00 | 0.00 | 0.00 | 0.00 | 0.00 |
| *Actinotalea* | 0.00 | 0.01 | 0.00 | 0.00 | 0.00 | 0.00 | 0.00 | 0.00 | 0.00 | 0.00 | 0.00 | 0.00 |
| *Aeromonas* | 0.00 | 0.00 | 0.00 | 0.00 | 0.00 | 0.00 | 0.01 | 0.00 | 0.00 | 0.00 | 0.00 | 0.00 |
| *Afipia* | 0.00 | 0.00 | 0.00 | 0.00 | 0.00 | 0.00 | 0.02 | 0.21 | 0.47 | 0.00 | 0.00 | 0.00 |
| AKYH478 unclassified | 0.01 | 0.02 | 0.00 | 0.00 | 0.00 | 0.00 | 0.00 | 0.00 | 0.00 | 0.00 | 0.00 | 0.00 |
| *Alcaligenaceae* unclassified | 0.01 | 0.00 | 0.10 | 0.02 | 0.00 | 0.08 | 0.00 | 0.05 | 0.10 | 0.07 | 0.19 | 0.25 |
| *Alphaproteobacteria* unclassified | 0.00 | 0.00 | 0.01 | 0.00 | 0.00 | 0.00 | 0.01 | 3.93 | 1.19 | 0.00 | 0.00 | 0.00 |
| *Anaerolineaceae* unclassified | 0.00 | 0.02 | 0.00 | 0.00 | 0.00 | 0.00 | 0.00 | 0.00 | 0.00 | 0.00 | 0.00 | 0.00 |
| *Bacillaceae* unclassified | 2.03 | 1.15 | 0.00 | 14.77 | 54.54 | 72.97 | 0.00 | 0.00 | 0.00 | 0.39 | 0.12 | 0.70 |
| *Bacillales* unclassified | 0.15 | 0.02 | 0.01 | 0.32 | 0.42 | 0.05 | 0.02 | 0.00 | 0.00 | 0.00 | 0.00 | 0.01 |
| *Bacilli* unclassified | 0.01 | 0.00 | 0.00 | 0.00 | 0.01 | 0.00 | 0.00 | 0.00 | 0.00 | 0.00 | 0.00 | 0.00 |
| *Bacillus* | 69.78 | 0.01 | 26.91 | 0.15 | 0.31 | 0.00 | 10.37 | 0.05 | 0.21 | 0.02 | 0.40 | 0.00 |
| *Bacteria* unclassified | 0.00 | 0.02 | 0.00 | 0.02 | 0.01 | 0.00 | 0.00 | 0.00 | 0.00 | 0.00 | 0.00 | 0.00 |
| *Bauldia* | 0.00 | 0.01 | 0.00 | 0.00 | 0.00 | 0.00 | 0.00 | 0.00 | 0.00 | 0.00 | 0.00 | 0.00 |
| *Betaproteobacteria* unclassified | 0.00 | 0.00 | 0.02 | 0.00 | 0.00 | 0.01 | 0.00 | 0.00 | 0.00 | 0.00 | 0.04 | 0.00 |
| *Blastocatella* | 0.00 | 0.01 | 0.00 | 0.00 | 0.00 | 0.00 | 0.00 | 0.00 | 0.00 | 0.00 | 0.00 | 0.00 |
| *Bordetella* | 0.06 | 0.11 | 0.47 | 0.20 | 0.03 | 2.10 | 0.22 | 1.51 | 4.78 | 8.08 | 42.89 | 25.40 |
| *Bosea* | 0.00 | 0.01 | 0.14 | 0.01 | 0.00 | 0.70 | 0.00 | 0.11 | 0.06 | 0.49 | 0.26 | 1.41 |
| *Bradyrhizobiaceae* unclassified | 0.00 | 0.00 | 0.00 | 0.01 | 0.00 | 0.00 | 0.00 | 0.00 | 0.00 | 0.02 | 0.01 | 0.02 |
| *Bradyrhizobium* | 0.00 | 0.02 | 0.00 | 0.00 | 0.00 | 0.01 | 0.09 | 0.06 | 0.33 | 0.59 | 0.12 | 0.23 |
| *Brevibacillus* | 14.78 | 1.21 | 2.41 | 3.82 | 5.23 | 1.29 | 1.18 | 0.02 | 0.03 | 0.14 | 1.13 | 0.02 |
| *Burkholderiaceae* unclassified | 0.00 | 0.00 | 0.00 | 0.00 | 0.00 | 0.00 | 0.00 | 0.00 | 0.02 | 0.00 | 0.00 | 0.00 |
| *Burkholderiales* unclassified | 0.00 | 0.00 | 0.00 | 0.00 | 0.00 | 0.00 | 0.00 | 0.00 | 0.00 | 0.01 | 0.00 | 0.00 |
| *Candidate division* OD1 unclassified | 0.00 | 0.00 | 0.00 | 0.00 | 0.00 | 0.00 | 0.00 | 0.00 | 0.01 | 0.00 | 0.00 | 0.00 |
| *Candidatus Planktomarina* DC5-80-3 | 0.00 | 0.00 | 0.00 | 0.05 | 0.00 | 0.00 | 0.00 | 0.00 | 0.00 | 0.00 | 0.00 | 0.00 |

| | | | | | | | | | | | | |
|---|---|---|---|---|---|---|---|---|---|---|---|---|
| *Castellaniella* | 3.39 | 0.14 | 18.26 | 4.28 | 1.44 | 2.28 | 5.80 | 0.03 | 5.25 | 2.87 | 0.85 | 3.08 |
| *Chelatococcus* | 0.00 | 0.00 | 0.00 | 0.00 | 0.00 | 0.00 | 0.00 | 0.22 | 0.06 | 0.00 | 0.00 | 0.00 |
| *Chitinophagaceae* unclassified | 0.00 | 0.00 | 0.00 | 0.00 | 0.00 | 0.00 | 0.01 | 0.00 | 0.00 | 0.01 | 0.01 | 0.01 |
| *Clostridiaceae* 1 unclassified | 0.00 | 0.00 | 0.00 | 0.62 | 0.02 | 0.01 | 0.00 | 0.00 | 0.00 | 0.00 | 0.00 | 0.00 |
| *Clostridium sensu stricto* 1 | 0.00 | 0.00 | 0.00 | 4.10 | 0.11 | 0.13 | 0.00 | 0.00 | 0.00 | 0.05 | 0.00 | 0.00 |
| *Clostridium sensu stricto* 3 | 0.10 | 0.00 | 1.50 | 0.48 | 0.46 | 0.38 | 0.13 | 0.00 | 0.14 | 0.07 | 0.06 | 0.00 |
| *Clostridium sensu stricto* 10 | 0.00 | 0.00 | 0.00 | 6.61 | 0.00 | 0.33 | 0.00 | 0.00 | 0.00 | 0.00 | 0.00 | 0.00 |
| *Clostridium sensu stricto* 11 | 0.00 | 0.00 | 0.00 | 0.00 | 0.00 | 0.00 | 0.00 | 0.00 | 0.01 | 0.00 | 0.00 | 0.00 |
| *Clostridium sensu stricto* 13 | 0.00 | 0.00 | 0.00 | 0.09 | 0.00 | 0.00 | 0.00 | 0.00 | 0.00 | 0.00 | 0.00 | 0.00 |
| *Comamonadaceae* unclassified | 0.04 | 0.00 | 0.01 | 0.01 | 0.00 | 0.00 | 0.01 | 0.04 | 0.01 | 0.00 | 0.03 | 0.02 |
| *Devosia* | 0.00 | 0.03 | 0.00 | 0.08 | 0.00 | 0.82 | 0.12 | 0.04 | 0.55 | 8.23 | 0.03 | 5.24 |
| *Dokdonella* | 0.00 | 0.02 | 0.00 | 0.01 | 0.00 | 0.00 | 0.15 | 0.08 | 0.00 | 0.00 | 0.05 | 0.00 |
| *Ensifer* | 0.00 | 0.25 | 0.01 | 0.00 | 0.00 | 0.00 | 0.04 | 1.79 | 0.31 | 0.00 | 0.00 | 0.01 |
| *Escherichia-Shigella* | 0.00 | 0.08 | 0.01 | 0.00 | 0.00 | 0.00 | 0.00 | 0.00 | 0.03 | 0.05 | 0.01 | 0.01 |
| *Family I* unclassified | 0.00 | 0.00 | 0.00 | 0.01 | 0.00 | 0.00 | 0.00 | 0.00 | 0.00 | 0.00 | 0.00 | 0.00 |
| *Ferrovibrio* | 0.00 | 0.08 | 0.00 | 0.00 | 0.00 | 0.00 | 0.00 | 0.13 | 0.01 | 0.00 | 0.00 | 0.00 |
| *Filimonas* | 0.00 | 0.00 | 0.00 | 0.00 | 0.00 | 0.00 | 0.04 | 0.00 | 0.03 | 0.00 | 0.00 | 0.00 |
| *Flavobacterium* | 0.00 | 0.00 | 0.00 | 0.00 | 0.00 | 0.00 | 0.00 | 0.00 | 0.00 | 0.02 | 0.00 | 0.00 |
| *Fluviicola* | 0.00 | 0.02 | 0.00 | 0.00 | 0.00 | 0.00 | 0.00 | 0.00 | 0.00 | 0.00 | 0.00 | 0.00 |
| *Fontibacillus* | 0.00 | 0.00 | 0.00 | 0.00 | 0.00 | 0.00 | 0.00 | 0.00 | 0.00 | 0.00 | 0.01 | 0.00 |
| *Gammaproteobacteria* unclassified | 0.00 | 0.01 | 0.00 | 0.01 | 0.00 | 0.00 | 0.00 | 0.00 | 0.00 | 0.00 | 0.00 | 0.00 |
| *Gordonia* | 0.00 | 1.89 | 0.02 | 0.01 | 0.01 | 0.26 | 0.01 | 0.50 | 0.13 | 0.76 | 0.28 | 1.15 |
| *Hydrogenophaga* | 7.49 | 72.96 | 24.17 | 24.41 | 0.19 | 6.31 | 30.60 | 50.72 | 37.19 | 29.06 | 30.17 | 24.57 |
| *Hyphomicrobium* | 0.00 | 0.00 | 0.00 | 0.00 | 0.00 | 0.00 | 0.06 | 0.10 | 0.16 | 0.04 | 0.00 | 0.00 |
| *Inquilinus* | 0.00 | 0.00 | 0.00 | 0.00 | 0.01 | 0.00 | 0.00 | 0.00 | 0.00 | 0.00 | 0.00 | 0.00 |
| *Kaistia* | 0.00 | 0.00 | 0.00 | 0.00 | 0.00 | 0.01 | 0.00 | 0.03 | 0.00 | 0.00 | 0.00 | 0.01 |
| KCM-B-112 unclassified | 0.00 | 0.01 | 0.00 | 0.00 | 0.00 | 0.00 | 0.00 | 0.00 | 0.00 | 0.00 | 0.00 | 0.00 |
| *Luminiphilus* | 0.00 | 0.00 | 0.00 | 0.01 | 0.00 | 0.00 | 0.00 | 0.00 | 0.00 | 0.00 | 0.00 | 0.00 |
| *Lysinibacillus* | 0.89 | 0.01 | 0.33 | 0.53 | 1.35 | 0.20 | 0.03 | 0.00 | 0.01 | 0.12 | 0.68 | 0.02 |
| *Microvirga* | 0.00 | 0.04 | 0.08 | 0.04 | 0.00 | 0.33 | 0.17 | 0.03 | 0.01 | 0.33 | 0.14 | 0.00 |
| MNC12 unclassified | 0.00 | 0.02 | 0.00 | 0.00 | 0.00 | 0.00 | 0.00 | 0.00 | 0.00 | 0.00 | 0.00 | 0.00 |
| *Moraxellaceae* unclassified | 0.00 | 0.01 | 0.00 | 0.00 | 0.00 | 0.00 | 0.00 | 0.00 | 0.00 | 0.00 | 0.00 | 0.00 |

| | | | | | | | | | | | | |
|---|---|---|---|---|---|---|---|---|---|---|---|---|
| *Mycobacterium* | 0.00 | 0.00 | 0.00 | 0.00 | 0.01 | 0.00 | 0.00 | 0.00 | 0.00 | 0.00 | 0.00 | 0.00 |
| *Mycoplana* | 0.11 | 0.28 | 0.17 | 1.00 | 0.00 | 0.24 | 0.05 | 0.14 | 0.01 | 0.10 | 0.08 | 0.04 |
| *Nitrosomonas* | 0.00 | 0.00 | 0.01 | 0.00 | 0.00 | 0.00 | 0.00 | 0.00 | 0.00 | 0.00 | 0.00 | 0.00 |
| *Nitrospira* | 0.00 | 0.01 | 0.00 | 0.00 | 0.00 | 0.00 | 0.00 | 0.00 | 0.00 | 0.00 | 0.00 | 0.00 |
| *Nocardia* | 0.00 | 0.04 | 0.00 | 0.00 | 0.00 | 0.00 | 0.00 | 0.02 | 0.06 | 0.01 | 0.17 | 0.02 |
| *Nocardiaceae* unclassified | 0.00 | 0.01 | 0.00 | 0.00 | 0.00 | 0.00 | 0.00 | 0.00 | 0.00 | 0.00 | 0.00 | 0.00 |
| *Nocardioidaceae* unclassified | 0.00 | 0.01 | 0.00 | 0.00 | 0.00 | 0.00 | 0.00 | 0.01 | 0.00 | 0.00 | 0.00 | 0.00 |
| *Nocardioides* | 0.59 | 4.21 | 0.58 | 0.00 | 0.00 | 0.04 | 0.24 | 1.42 | 0.67 | 0.02 | 2.12 | 0.05 |
| *Nordella* | 0.00 | 0.00 | 0.00 | 0.00 | 0.00 | 0.01 | 0.01 | 0.01 | 0.08 | 0.08 | 0.05 | 0.20 |
| *Novosphingobium* | 0.00 | 0.01 | 0.00 | 0.00 | 0.00 | 0.00 | 0.00 | 0.00 | 0.00 | 0.00 | 0.00 | 0.00 |
| *Obscuribacterales* unclassified | 0.00 | 0.01 | 0.00 | 0.00 | 0.00 | 0.00 | 0.00 | 0.00 | 0.00 | 0.00 | 0.00 | 0.00 |
| OPB56 unclassified | 0.00 | 0.01 | 0.00 | 0.00 | 0.00 | 0.01 | 1.26 | 2.05 | 3.04 | 0.00 | 0.00 | 0.03 |
| *Opitutus* | 0.00 | 0.00 | 0.00 | 0.00 | 0.00 | 0.00 | 0.00 | 0.00 | 0.01 | 0.38 | 0.13 | 1.06 |
| *Paenibacillaceae* unclassified | 0.00 | 0.00 | 0.00 | 0.06 | 0.03 | 0.03 | 0.00 | 0.00 | 0.00 | 0.00 | 0.03 | 0.00 |
| *Paenibacillus* | 0.00 | 0.18 | 0.00 | 17.36 | 32.49 | 2.15 | 0.00 | 0.07 | 0.00 | 0.37 | 0.64 | 0.08 |
| *Parvibaculum* | 0.01 | 0.09 | 0.00 | 0.00 | 0.00 | 0.00 | 0.25 | 0.86 | 1.24 | 1.33 | 0.01 | 3.95 |
| *Perlucidibaca* | 0.00 | 0.02 | 0.00 | 0.11 | 0.00 | 0.01 | 10.11 | 11.99 | 1.00 | 0.17 | 0.00 | 0.05 |
| *Phenylobacterium* | 0.00 | 0.00 | 0.00 | 0.02 | 0.00 | 0.10 | 0.00 | 0.00 | 0.00 | 0.10 | 0.00 | 0.33 |
| *Phyllobacteriaceae* unclassified | 0.05 | 0.93 | 0.23 | 2.12 | 0.05 | 0.61 | 1.22 | 1.26 | 11.02 | 1.93 | 4.21 | 6.67 |
| *Pibocella* | 0.00 | 0.00 | 0.00 | 0.01 | 0.00 | 0.00 | 0.00 | 0.00 | 0.00 | 0.00 | 0.00 | 0.00 |
| *Planctomyces* | 0.00 | 0.02 | 0.00 | 0.00 | 0.00 | 0.00 | 0.00 | 0.00 | 0.00 | 0.00 | 0.00 | 0.00 |
| *Planococcaceae* unclassified | 0.02 | 0.00 | 0.01 | 15.62 | 3.20 | 1.88 | 0.00 | 0.00 | 0.00 | 0.06 | 0.01 | 0.11 |
| *Pseudaminobacter* | 0.04 | 0.69 | 0.00 | 0.00 | 0.00 | 0.00 | 0.02 | 0.08 | 0.00 | 0.00 | 0.00 | 0.00 |
| *Pseudolabrys* | 0.00 | 0.00 | 0.00 | 0.01 | 0.00 | 0.00 | 0.00 | 0.00 | 0.02 | 0.01 | 0.01 | 0.00 |
| *Pseudomonas* | 0.00 | 0.01 | 0.00 | 0.00 | 0.00 | 0.00 | 0.00 | 0.00 | 0.00 | 0.00 | 0.00 | 0.00 |
| *Pseudoxanthomonas* | 0.02 | 0.06 | 9.46 | 0.04 | 0.00 | 0.06 | 4.96 | 8.64 | 13.72 | 0.05 | 0.20 | 0.03 |
| *Ramlibacter* | 0.00 | 0.07 | 0.00 | 0.00 | 0.00 | 0.01 | 0.00 | 0.10 | 0.02 | 0.00 | 0.01 | 0.06 |
| *Reyranella* | 0.00 | 0.00 | 0.00 | 0.00 | 0.00 | 0.00 | 0.00 | 0.22 | 0.05 | 0.00 | 0.00 | 0.00 |
| *Rheinheimera* | 0.00 | 0.01 | 0.00 | 0.00 | 0.00 | 0.00 | 0.00 | 0.00 | 0.01 | 0.00 | 0.00 | 0.00 |
| *Rhizobiaceae* unclassified | 0.00 | 0.01 | 0.00 | 0.00 | 0.00 | 0.00 | 0.00 | 0.00 | 0.00 | 0.00 | 0.00 | 0.00 |
| *Rhizobiales* unclassified | 0.00 | 0.00 | 0.00 | 0.01 | 0.00 | 0.01 | 0.01 | 0.06 | 0.07 | 0.06 | 0.01 | 0.12 |
| *Rhizobium* | 0.00 | 0.01 | 0.00 | 0.00 | 0.00 | 0.01 | 0.00 | 0.07 | 0.07 | 0.00 | 0.00 | 0.00 |

| | | | | | | | | | | | | |
|---|---|---|---|---|---|---|---|---|---|---|---|---|
| *Rhodobacteraceae* unclassified | 0.00 | 0.00 | 0.00 | 0.01 | 0.00 | 0.00 | 0.00 | 0.03 | 0.00 | 0.00 | 0.00 | 0.00 |
| *Rhodobium* | 0.00 | 0.00 | 0.00 | 0.01 | 0.00 | 0.00 | 0.00 | 0.00 | 0.00 | 0.00 | 0.00 | 0.00 |
| *Rhodococcus* | 0.00 | 11.17 | 0.06 | 0.01 | 0.01 | 0.10 | 0.08 | 2.39 | 1.96 | 0.19 | 0.25 | 2.14 |
| *Rhodospirillaceae* unclassified | 0.00 | 0.01 | 0.00 | 0.00 | 0.00 | 0.00 | 0.00 | 0.00 | 0.00 | 0.00 | 0.00 | 0.00 |
| *Rhodospirillales* unclassified | 0.00 | 0.01 | 0.00 | 0.00 | 0.00 | 0.00 | 0.00 | 0.00 | 0.00 | 0.00 | 0.00 | 0.00 |
| *Schlegelella* | 0.00 | 0.00 | 0.01 | 0.00 | 0.00 | 0.00 | 0.00 | 0.00 | 0.00 | 0.00 | 0.00 | 0.00 |
| *Shigella* | 0.00 | 0.00 | 0.00 | 0.00 | 0.00 | 0.01 | 0.00 | 0.14 | 0.33 | 0.00 | 0.00 | 0.00 |
| *Sphingomonadaceae* unclassified | 0.00 | 0.00 | 0.01 | 0.00 | 0.00 | 0.00 | 0.00 | 0.00 | 0.00 | 0.00 | 0.00 | 0.00 |
| *Sphingomonadales* unclassified | 0.01 | 0.00 | 0.00 | 0.01 | 0.00 | 0.00 | 0.01 | 0.00 | 0.00 | 0.01 | 0.00 | 0.01 |
| *Sphingomonas* | 0.02 | 0.17 | 0.01 | 0.00 | 0.00 | 0.00 | 0.02 | 0.02 | 0.00 | 0.00 | 0.00 | 0.00 |
| *Sphingopyxis* | 0.39 | 3.66 | 6.17 | 2.85 | 0.06 | 5.60 | 26.46 | 10.37 | 12.86 | 39.46 | 5.92 | 19.04 |
| *Streptomyces* | 0.00 | 0.01 | 0.00 | 0.00 | 0.01 | 0.00 | 0.00 | 0.00 | 0.00 | 0.00 | 0.00 | 0.00 |
| *Subgroup 6* unclassified | 0.00 | 0.01 | 0.00 | 0.00 | 0.00 | 0.00 | 0.00 | 0.00 | 0.00 | 0.00 | 0.00 | 0.00 |
| *Sulfitobacter* | 0.00 | 0.00 | 0.00 | 0.01 | 0.00 | 0.00 | 0.00 | 0.00 | 0.00 | 0.00 | 0.00 | 0.00 |
| SV1-3 | 0.00 | 0.00 | 0.00 | 0.01 | 0.00 | 0.00 | 0.00 | 0.00 | 0.00 | 0.00 | 0.00 | 0.00 |
| Sva0071 unclassified | 0.00 | 0.00 | 0.00 | 0.01 | 0.00 | 0.00 | 0.00 | 0.00 | 0.00 | 0.00 | 0.00 | 0.00 |
| *Terrimonas* | 0.00 | 0.03 | 8.82 | 0.04 | 0.00 | 0.85 | 6.20 | 0.31 | 2.64 | 4.25 | 8.67 | 3.85 |
| *Thermomonas* | 0.00 | 0.00 | 0.00 | 0.00 | 0.00 | 0.00 | 0.01 | 0.09 | 0.01 | 0.00 | 0.00 | 0.00 |
| *WD2101 soil group* unclassified | 0.00 | 0.01 | 0.00 | 0.00 | 0.00 | 0.00 | 0.00 | 0.00 | 0.00 | 0.00 | 0.00 | 0.00 |
| *Winogradskyella* | 0.00 | 0.00 | 0.00 | 0.01 | 0.00 | 0.00 | 0.00 | 0.00 | 0.00 | 0.00 | 0.00 | 0.00 |
| **Total Genera Counts** | **23** | **61** | **29** | **47** | **23** | **37** | **37** | **43** | **47** | **39** | **38** | **37** |

*Genera counts include any genus with a relative abundance above zero in a sample.

**SI Table S3.** Relative genera abundance and number of genera identified in samples from each experiment (Exp1 and Exp2) of consortia E1 and E2 at Day 0 and Day 14.

| | **Day 0** | | | | **Day 14** | | | |
|---|---|---|---|---|---|---|---|---|
| **sample** | **E1 Exp** | **E1 Exp2** | **E2 Exp1** | **E2 Exp2** | **E1 Exp** | **E1 Exp2** | **E2 Exp1** | **E2 Exp2** |
| 0319-6G20 unclassified | 0.00 | 0.00 | 0.00 | 0.00 | 0.02 | 0.00 | 0.00 | 0.00 |
| *Acetobacteraceae* unclassified | 0.00 | 0.00 | 0.00 | 0.00 | 0.00 | 0.00 | 0.02 | 0.00 |
| *Acidimicrobiaceae* unclassified | 0.01 | 0.00 | 0.00 | 0.00 | 0.00 | 0.00 | 0.00 | 0.00 |
| *Actinotalea* | 0.00 | 0.02 | 0.01 | 0.00 | 0.00 | 0.00 | 0.00 | 0.00 |

| | | | | | | | | |
|---|---|---|---|---|---|---|---|---|
| *Afipia* | 0.00 | 0.29 | 0.00 | 1.97 | 0.00 | 0.00 | 0.00 | 0.00 |
| *Agromyces* | 0.00 | 0.00 | 0.25 | 0.06 | 0.00 | 0.00 | 0.00 | 0.00 |
| AKYG1722 unclassified | 0.00 | 0.00 | 0.00 | 0.00 | 0.01 | 0.00 | 0.00 | 0.00 |
| AKYH478 unclassified | 0.00 | 0.01 | 0.00 | 0.00 | 0.00 | 0.00 | 0.00 | 0.00 |
| *Alphaproteobacteria* unclassified | 0.90 | 1.50 | 3.23 | 2.36 | 0.00 | 0.00 | 0.30 | 0.00 |
| *Arenimonas* | 0.00 | 0.00 | 0.00 | 0.00 | 0.00 | 0.00 | 0.11 | 0.00 |
| *Bacillus* | 0.02 | 0.00 | 0.00 | 0.00 | 0.00 | 0.00 | 0.20 | 0.00 |
| *Bacteria* unclassified | 0.04 | 0.00 | 0.05 | 0.00 | 0.22 | 0.00 | 0.02 | 0.00 |
| *Bauldia* | 0.02 | 0.00 | 0.05 | 0.00 | 0.04 | 0.06 | 0.00 | 0.01 |
| *Beijerinckiaceae* unclassified | 0.00 | 0.01 | 0.00 | 0.00 | 0.00 | 0.00 | 0.00 | 0.00 |
| *Betaproteobacteria* unclassified | 0.00 | 0.01 | 0.00 | 0.00 | 0.27 | 0.06 | 0.02 | 0.00 |
| *Blastocatella* | 0.00 | 0.00 | 0.00 | 0.00 | 0.41 | 1.99 | 0.00 | 0.14 |
| *Bordetella* | 0.00 | 0.00 | 0.15 | 0.00 | 0.94 | 0.05 | 0.01 | 0.00 |
| *Bosea* | 0.00 | 0.00 | 0.00 | 0.17 | 0.01 | 0.02 | 0.00 | 0.00 |
| *Bradyrhizobiaceae* unclassified | 0.00 | 0.00 | 0.00 | 0.00 | 0.04 | 0.00 | 0.00 | 0.07 |
| *Bradyrhizobium* | 0.00 | 0.09 | 0.37 | 0.30 | 0.06 | 0.14 | 0.18 | 1.86 |
| *Brevibacillus* | 0.03 | 0.02 | 0.00 | 0.00 | 0.06 | 0.00 | 0.00 | 0.00 |
| *Burkholderiaceae* unclassified | 0.00 | 0.00 | 0.00 | 0.01 | 0.00 | 0.00 | 0.00 | 0.00 |
| *Candidatus Odyssella* | 0.00 | 0.00 | 0.00 | 0.00 | 0.00 | 0.00 | 0.03 | 0.00 |
| *Castellaniella* | 0.00 | 0.00 | 0.00 | 0.00 | 0.00 | 0.00 | 0.16 | 0.00 |
| *Chelatococcus* | 0.05 | 0.24 | 1.68 | 0.30 | 0.01 | 0.00 | 0.00 | 0.00 |
| *Chitinophagaceae* unclassified | 0.01 | 0.00 | 0.00 | 0.00 | 0.00 | 0.00 | 0.00 | 0.00 |
| *Chthoniobacterales* unclassified | 0.35 | 0.11 | 0.08 | 0.25 | 0.00 | 0.00 | 0.00 | 0.00 |
| *Clostridium sensu stricto 12* | 0.00 | 0.00 | 0.00 | 0.00 | 0.00 | 0.01 | 0.00 | 0.00 |
| *Comamonadaceae* unclassified | 0.01 | 0.10 | 0.00 | 0.01 | 0.00 | 0.06 | 0.00 | 0.00 |
| *Corynebacterium* | 0.00 | 0.00 | 0.00 | 0.00 | 0.00 | 0.00 | 0.00 | 0.01 |
| ctg-CGOF202 unclassified | 0.00 | 0.01 | 0.00 | 0.00 | 0.00 | 0.00 | 0.00 | 0.00 |
| *Devosia* | 0.00 | 0.00 | 0.02 | 0.02 | 2.25 | 3.22 | 7.65 | 6.77 |
| *Escherichia-Shigella* | 0.04 | 0.00 | 0.01 | 0.00 | 0.02 | 0.00 | 0.01 | 0.00 |
| *Flavobacterium* | 0.00 | 0.00 | 0.00 | 0.00 | 0.00 | 0.00 | 0.00 | 0.01 |
| *Gaiellales* unclassified | 0.00 | 0.02 | 0.00 | 0.00 | 0.02 | 0.01 | 0.00 | 0.00 |

| | | | | | | | | |
|---|---|---|---|---|---|---|---|---|
| *Gemmatimonadaceae* unclassified | 0.00 | 0.00 | 0.00 | 0.00 | 3.84 | 3.11 | 0.06 | 0.05 |
| *Gordonia* | 0.03 | 0.02 | 0.02 | 0.01 | 0.00 | 0.00 | 0.00 | 0.00 |
| *Hydrogenophaga* | 66.02 | 54.88 | 32.77 | 14.18 | 70.04 | 51.11 | 40.56 | 9.48 |
| *Hyphomicrobiaceae* unclassified | 0.00 | 0.01 | 0.00 | 0.02 | 0.01 | 0.02 | 0.00 | 0.00 |
| *Hyphomicrobium* | 0.01 | 0.54 | 1.87 | 0.54 | 0.01 | 0.77 | 0.52 | 0.73 |
| *Iamia* | 0.13 | 0.02 | 0.01 | 0.00 | 0.11 | 0.01 | 0.00 | 0.00 |
| JG30-KF-CM45 unclassified | 0.00 | 0.04 | 0.01 | 0.02 | 0.00 | 0.00 | 0.00 | 0.02 |
| JG30-KF-CM66 unclassified | 0.00 | 0.01 | 0.00 | 0.00 | 0.00 | 0.01 | 0.00 | 0.00 |
| *Legionella* | 0.00 | 0.00 | 0.11 | 0.05 | 0.00 | 0.00 | 0.00 | 0.00 |
| *Limnobacter* | 0.12 | 0.23 | 0.00 | 0.04 | 0.00 | 0.00 | 0.00 | 0.00 |
| *Lysinimonas* | 0.00 | 0.01 | 0.00 | 0.00 | 0.00 | 0.00 | 0.00 | 0.00 |
| *Microvirga* | 0.01 | 0.02 | 0.07 | 0.00 | 0.04 | 0.14 | 1.39 | 1.79 |
| *Mycobacterium* | 0.06 | 0.33 | 0.00 | 0.00 | 0.00 | 0.00 | 0.00 | 0.00 |
| *Mycoplana* | 0.00 | 0.02 | 0.00 | 0.02 | 0.01 | 0.01 | 0.00 | 0.00 |
| *Nitratireductor* | 0.00 | 0.00 | 0.00 | 0.00 | 0.00 | 0.00 | 0.01 | 0.00 |
| *Nocardia* | 0.03 | 0.09 | 0.01 | 0.23 | 0.01 | 0.04 | 0.02 | 0.04 |
| *Nocardiaceae* unclassified | 0.01 | 0.00 | 0.00 | 0.01 | 0.00 | 0.01 | 0.00 | 0.00 |
| *Nocardioides* | 0.00 | 0.00 | 0.00 | 0.00 | 0.00 | 0.00 | 0.04 | 0.00 |
| *Nordella* | 0.00 | 0.10 | 0.05 | 0.12 | 0.06 | 0.11 | 0.12 | 0.26 |
| OPB35 soil group unclassified | 0.01 | 0.00 | 0.00 | 0.00 | 0.00 | 0.00 | 0.00 | 0.00 |
| OPB56 unclassified | 0.00 | 0.00 | 0.00 | 0.00 | 0.00 | 0.00 | 0.02 | 0.00 |
| *Opitutaceae* unclassified | 0.00 | 0.02 | 0.00 | 0.00 | 0.00 | 0.00 | 0.00 | 0.00 |
| *Opitutus* | 16.53 | 7.75 | 0.40 | 2.50 | 0.02 | 0.03 | 0.00 | 0.01 |
| *Paenibacillaceae* unclassified | 0.00 | 0.00 | 0.00 | 0.00 | 0.00 | 0.00 | 0.00 | 0.00 |
| *Parvibaculum* | 0.19 | 0.23 | 5.05 | 0.22 | 0.03 | 0.00 | 0.17 | 0.01 |
| *Pedomicrobium* | 0.02 | 0.09 | 0.21 | 0.06 | 0.14 | 0.05 | 0.16 | 0.19 |
| *Perlucidibaca* | 0.01 | 0.00 | 0.00 | 0.00 | 0.01 | 0.00 | 0.04 | 0.00 |
| *Phenylobacterium* | 0.00 | 0.00 | 0.00 | 0.00 | 0.00 | 0.00 | 0.00 | 0.01 |
| *Phyllobacteriaceae* unclassified | 0.25 | 0.22 | 7.87 | 18.28 | 0.50 | 0.24 | 17.13 | 13.35 |
| *Proteobacteria* unclassified | 0.01 | 0.00 | 0.00 | 0.00 | 0.00 | 0.00 | 0.00 | 0.01 |
| *Pseudolabrys* | 0.00 | 0.22 | 0.00 | 0.04 | 0.19 | 0.03 | 0.02 | 0.08 |

| | | | | | | | | |
|---|---|---|---|---|---|---|---|---|
| *Pseudoxanthomonas* | 0.70 | 1.54 | 7.32 | 6.52 | 0.00 | 0.00 | 0.08 | 0.01 |
| *Ramlibacter* | 0.00 | 0.04 | 0.00 | 0.26 | 0.01 | 0.03 | 0.00 | 0.29 |
| *Reyranella* | 0.50 | 0.48 | 0.68 | 0.39 | 0.28 | 0.30 | 3.70 | 0.48 |
| *Rhizobiales* unclassified | 0.26 | 0.77 | 0.08 | 0.08 | 2.81 | 2.52 | 0.61 | 0.26 |
| *Rhizobium* | 0.00 | 0.03 | 0.00 | 0.03 | 0.00 | 0.02 | 0.00 | 0.00 |
| *Rhodococcus* | 8.67 | 3.12 | 7.68 | 33.11 | 12.08 | 18.79 | 2.92 | 35.67 |
| *Rhodoplanes* | 0.08 | 0.15 | 0.02 | 0.03 | 0.09 | 0.04 | 0.04 | 0.03 |
| *Schlegelella* | 0.00 | 0.00 | 0.00 | 0.00 | 0.01 | 0.00 | 0.00 | 0.00 |
| SJA-149 unclassified | 0.10 | 0.11 | 0.14 | 0.29 | 0.32 | 0.09 | 0.13 | 0.23 |
| *Sphingomonadaceae* unclassified | 0.00 | 0.00 | 0.00 | 0.01 | 0.00 | 0.00 | 0.02 | 0.04 |
| *Sphingomonadales* unclassified | 0.00 | 0.07 | 0.01 | 0.00 | 0.00 | 0.00 | 0.00 | 0.01 |
| *Sphingomonas* | 0.00 | 0.00 | 0.00 | 0.01 | 0.07 | 0.00 | 0.05 | 0.02 |
| *Sphingopyxis* | 0.65 | 4.99 | 29.15 | 4.19 | 3.76 | 9.38 | 23.37 | 18.70 |
| *Stella* | 0.00 | 0.00 | 0.00 | 0.00 | 0.01 | 0.00 | 0.01 | 0.00 |
| *Stenotrophomonas* | 0.00 | 0.01 | 0.00 | 0.01 | 0.00 | 0.00 | 0.00 | 0.00 |
| *Terrimonas* | 4.06 | 21.40 | 0.52 | 12.99 | 0.71 | 5.78 | 0.10 | 4.26 |
| *Thermomonas* | 0.00 | 0.01 | 0.00 | 0.28 | 0.42 | 1.74 | 0.00 | 5.09 |
| TK10 unclassified | 0.01 | 0.00 | 0.00 | 0.00 | 0.03 | 0.00 | 0.00 | 0.00 |
| *Truepera* | 0.05 | 0.00 | 0.05 | 0.00 | 0.00 | 0.00 | 0.00 | 0.00 |
| *Turneriella* | 0.00 | 0.00 | 0.00 | 0.00 | 0.00 | 0.00 | 0.00 | 0.01 |
| *Vibrio* | 0.00 | 0.00 | 0.00 | 0.01 | 0.00 | 0.00 | 0.00 | 0.00 |
| **Total Genera Counts** | **28** | **36** | **25** | **33** | **31** | **29** | **27** | **31** |

*Genera counts include any genus with a relative abundance above zero in a sample.
